# S:D614G and S:H655Y are gateway mutations that act epistatically to promote SARS-CoV-2 variant fitness

**DOI:** 10.1101/2023.03.30.535005

**Authors:** Leonid Yurkovetskiy, Shawn Egri, Chaitanya Kurhade, Marco A. Diaz-Salinas, Javier A. Jaimes, Thomas Nyalile, Xuping Xie, Manish C. Choudhary, Ann Dauphin, Jonathan Z. Li, James B. Munro, Pei-Yong Shi, Kuang Shen, Jeremy Luban

**Affiliations:** Program in Molecular Medicine, University of Massachusetts Medical School, Worcester, MA 01605, USA; Massachusetts Consortium on Pathogen Readiness, Boston, MA, 02115; Department of Biochemistry and Molecular Biology, University of Texas Medical Branch, Galveston, Texas, USA; Department of Microbiology and Physiological Systems, University of Massachusetts Medical School, Worcester, MA 01605, USA; Department of Medicine, Brigham and Women’s Hospital, Boston, MA 02115, USA; Department of Biochemistry and Molecular Biotechnology, University of Massachusetts Medical School, Worcester, MA 01605, USA; Broad Institute of Harvard and MIT, Cambridge, MA 02142, USA; Ragon Institute of MGH, MIT and Harvard, Cambridge, MA 02139, USA

**Keywords:** SARS-CoV-2, COVID-19, coronavirus, variants, Spike protein, epistasis, evolution, ACE2, pandemic, infectivity, S:Q613H, S:D614G, S:H655Y

## Abstract

SARS-CoV-2 variants bearing complex combinations of mutations that confer increased transmissibility, COVID-19 severity, and immune escape, were first detected after S:D614G had gone to fixation, and likely originated during persistent infection of immunocompromised hosts. To test the hypothesis that S:D614G facilitated emergence of such variants, S:D614G was reverted to the ancestral sequence in the context of sequential Spike sequences from an immunocompromised individual, and within each of the major SARS-CoV-2 variants of concern. In all cases, infectivity of the S:D614G revertants was severely compromised. The infectivity of atypical SARS-CoV-2 lineages that propagated in the absence of S:D614G was found to be dependent upon either S:Q613H or S:H655Y. Notably, Gamma and Omicron variants possess both S:D614G and S:H655Y, each of which contributed to infectivity of these variants. Among sarbecoviruses, S:Q613H, S:D614G, and S:H655Y are only detected in SARS-CoV-2, which is also distinguished by a polybasic S1/S2 cleavage site. Genetic and biochemical experiments here showed that S:Q613H, S:D614G, and S:H655Y each stabilize Spike on virions, and that they are dispensable in the absence of S1/S2 cleavage, consistent with selection of these mutations by the S1/S2 cleavage site. CryoEM revealed that either S:D614G or S:H655Y shift the Spike receptor binding domain (RBD) towards the open conformation required for ACE2-binding and therefore on pathway for infection. Consistent with this, an smFRET reporter for RBD conformation showed that both S:D614G and S:H655Y spontaneously adopt the conformation that ACE2 induces in the parental Spike. Data from these orthogonal experiments demonstrate that S:D614G and S:H655Y are convergent adaptations to the polybasic S1/S2 cleavage site which stabilize S1 on the virion in the open RBD conformation and act epistatically to promote the fitness of variants bearing complex combinations of clinically significant mutations.

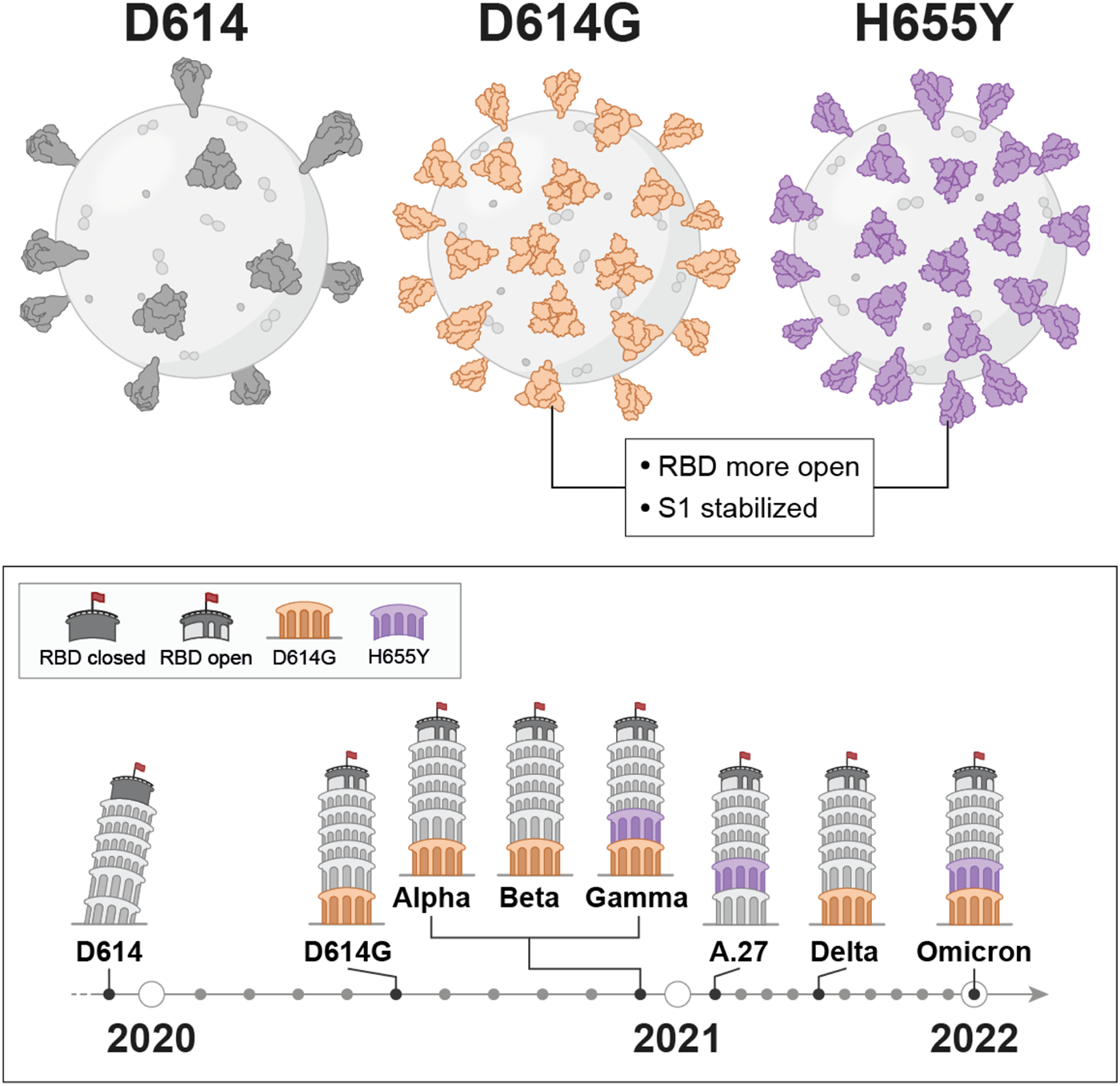

**Highlights:**

- S:D614G is ubiquitous among SARS-CoV-2 B-lineage Spikes and is required for infectivity of the main Variants of Concern
- In an example of convergent evolution, SARS-CoV-2 A lineage viruses maintained transmission chains in the absence of S:D614G, but were instead dependent upon S:Q613H or S:H655Y
- S:D614G and S:H655Y are both adaptations to the polybasic S1/S2 cleavage site
- Increased infectivity of S:D614G and S:H655Y is associated with a more open RBD conformation and increased steady-state levels of virion-associated S1

## Introduction

Severe acute respiratory syndrome coronavirus 2 (SARS-CoV-2), the cause of COVID-19^1–6^, is a member of the *Nidovirales*, an order that includes viruses with single-stranded, positive-sense, RNA genomes as large as 41 kb^7^. Among RNA viruses, *Nidovirales* are unique in that they encode a proofreading exoribonuclease thought necessary to prevent lethal mutagenesis of the large viral genome by the error-prone, viral RNA-dependent RNA polymerase^8–12^. The SARS-CoV-2 genome is 29.9 kb and indeed encodes a proof-reading exoribonuclease^13^. It was therefore not surprising that SARS-CoV-2 genomic RNA accumulated only two mutations per month over the first year of the pandemic^14–18^. Nonetheless, by September 2020, the S:D614G mutation constituted well over 98% of sequences submitted to GISAID on any given date, presumably due to its increased intrinsic infectivity and association with higher viral load and greater transmissibility^19–25^.

Towards the end of 2020, the SARS-CoV-2 pandemic was suddenly dominated by the Alpha, Beta, and Gamma Variants of Concern, each of which bore complex combinations of non-synonymous mutations that arose independently on three different continents. These variants were associated with greater transmissibility, increased pathogenicity, or decreased sensitivity to neutralization by Spike-specific antibodies^26–32^. Since then, the SARS-CoV-2 pandemic has been characterized by increasingly fit SARS-CoV-2 variants^33^, including Delta and its sublineages^34–36^, and Omicron, the latter having more than 30 nonsynonymous Spike mutations with respect to the ancestral virus^37^. Given the clinical and public health significance of the variants, it is critical that the conditions which selected large numbers of SARS-CoV-2 mutations at the end of 2020 be better understood.

One factor that changed in the first year of the pandemic was that SARS-CoV-2 initially infected an immunologically naive population, while in subsequent waves of infection the probability increased that the virus would encounter individuals with antiviral immunity elicited by prior infection or vaccination. Evidence for the significance of this change in selective pressure on SARS-CoV-2 comes from the increasing frequency over the course of the pandemic of mutations with decreased sensitivity to neutralizing antibodies^27–32, 37^.

Another possibility is that SARS-CoV-2 variants bearing complex sets of mutations accumulated at a steady rate within a context where viral sequences were not monitored^38^. In this scenario, complex variants would emerge fully-formed, without leaving a trail of precursor strains bearing intermediate numbers of mutations^18, 39, 40^. SARS-CoV-2 genomic surveillance has been minimal to nonexistent in much of the world, including in locations where retrospective serology indicated that the majority of the population had been infected with SARS-CoV-2^41^. One striking example of such emergence was provided by travelers from a country with no SARS-CoV-2 surveillance who were incidentally found to be infected with the highly unusual A.30 variant encoding a Spike with 16 amino acid changes^42^.

Reverse zoonosis is another possible source of SARS-CoV-2 variants with complex combinations of mutations. A number of animal species have been naturally-infected by SARS-CoV-2 and sustained virus transmission has been documented within mink and deer populations^43–45^. Ongoing transmission through animals would be expected to select for mutations adaptive to that species though the resulting SARS-CoV-2 strains might retain the ability to replicate in people with altered transmissibility, pathogenicity, or sensitivity to immune responses. Indeed, in reverse zoonoses from mink, hamster, and deer, people have been infected with SARS-CoV-2 strains not otherwise seen in humans^43, 44, 46, 47^.

Many mutations found in circulating SARS-CoV-2 variants have been detected in immunocompromised individuals with persistent infection^48–56^. Such individuals have thus been considered a likely source of the widely-circulating SARS-CoV-2 variants with complex sets of mutations^39^. Chronic, high-level SARS-CoV-2 replication within an immunocompromised host would provide the virus with time to explore genetic space in the absence of immune pressure, with greater probability of recombination as a result of two variants simultaneously infecting one cell, and without the bottleneck that comes with person-to-person transmission^39, 57–59^.

An increased number of SARS-CoV-2 mutations could be observed if the intrinsic error rate of the viral replication machinery increased. S:D614G was accompanied by a mutation in the viral RNA-dependent RNA polymerase, nsp12:P323L^20, 60^. In isolation, nsp12:P323L increased SARS-CoV-2 plaque size^60^, though the effect of this mutation on the replication error rate was not measured. In some persistent infections the rate of change in SARS-CoV-2 sequence was calculated to be increased^56, 61^, in other cases not^51^, and an increased rate of mutation was estimated in the branch of the phylogenetic tree that gave rise to the Alpha variant^18^.

Finally, the simultaneous appearance on three continents of different SARS-CoV-2 variants bearing complex sets of mutations, several months after the S:D614G mutation had gone to near fixation, suggests that extensive SARS-CoV-2 evolution after spillover from its animal reservoir was contingent upon prior establishment of S:D614G. This last hypothesis was tested here.

## Results

### SARS-CoV-2 Spike mutants that accumulate during prolonged infection are S:D614G-dependent

Sequential SARS-CoV-2 genomic sequences were reported from a person on immunosuppressive therapy for severe antiphospholipid syndrome who was continuously infected with SARS-CoV-2 for over five months^48^. The S amino acid sequence encoded by the earliest time-point had the S:D614G mutation (day 18 post-infection, Figure 1A) but was otherwise identical to that of the original SARS-CoV-2 Wuhan-Hu-1 (GenBank Accession NC_045512). By day 75, S coding sequence had gained four nonsynonymous mutations and a deletion of nine nucleotides (Figure 1A). On subsequent days an increasing number of nonsynonymous mutations was observed, including a number that were subsequently seen in Variants of Concern (Figure 1A).

To test the importance of S:D614G for the infectivity of the Spike proteins identified during this persistent infection, S sequences from each time-point were cloned into a mammalian expression plasmid, and, in each case, S:D614G was reverted back to S:D614. Infectivity was tested in a single-cycle, cell-free virion, infectivity assay using lentiviral pseudotypes, as previously described^20^. When S:D614G in the day 18 sequence was reverted back to the ancestral aspartic acid present in Wuhan-Hu-1, a 2.9-fold reduction in infectivity was observed (p=0.0001, red vs white bars, Figure 1B), as previously reported^20^. When sequences from later time-points bearing greater numbers of non-synonymous mutations were reverted back to the ancestral aspartic acid, greater decreases in infectivity were observed (Figure 1B), indicating even greater dependence of infectivity on S:D614G: day 75, 3.9-fold reduction (p=0.004); day 128, 16.2-fold reduction (p<0.0001); day 143, 14.9-fold reduction (p=0.0003); day 146, 19.4-fold reduction (p<0.0001).

Spike protein from each of the time-points was examined in western blots using antibodies specific for the S1 and S2 proteins. In cell lysate, reversion of the S:D614G mutation had minimal effect on Spike protein from any of the time-points (Figure 1C). In contrast, reversion of the S:D614G mutation reduced virion-associated S1 and S2 protein on the virus-like particles (Figure 1C). The magnitude was generally greater for S1 than for S2, and was greater at later time points (Figure 1C). These results demonstrate that, as greater numbers of non-synonymous mutations accumulate at later time points, S:D614G becomes increasingly important to maintain infectivity and Spike protein levels associated with virions.

**Figure 1.**
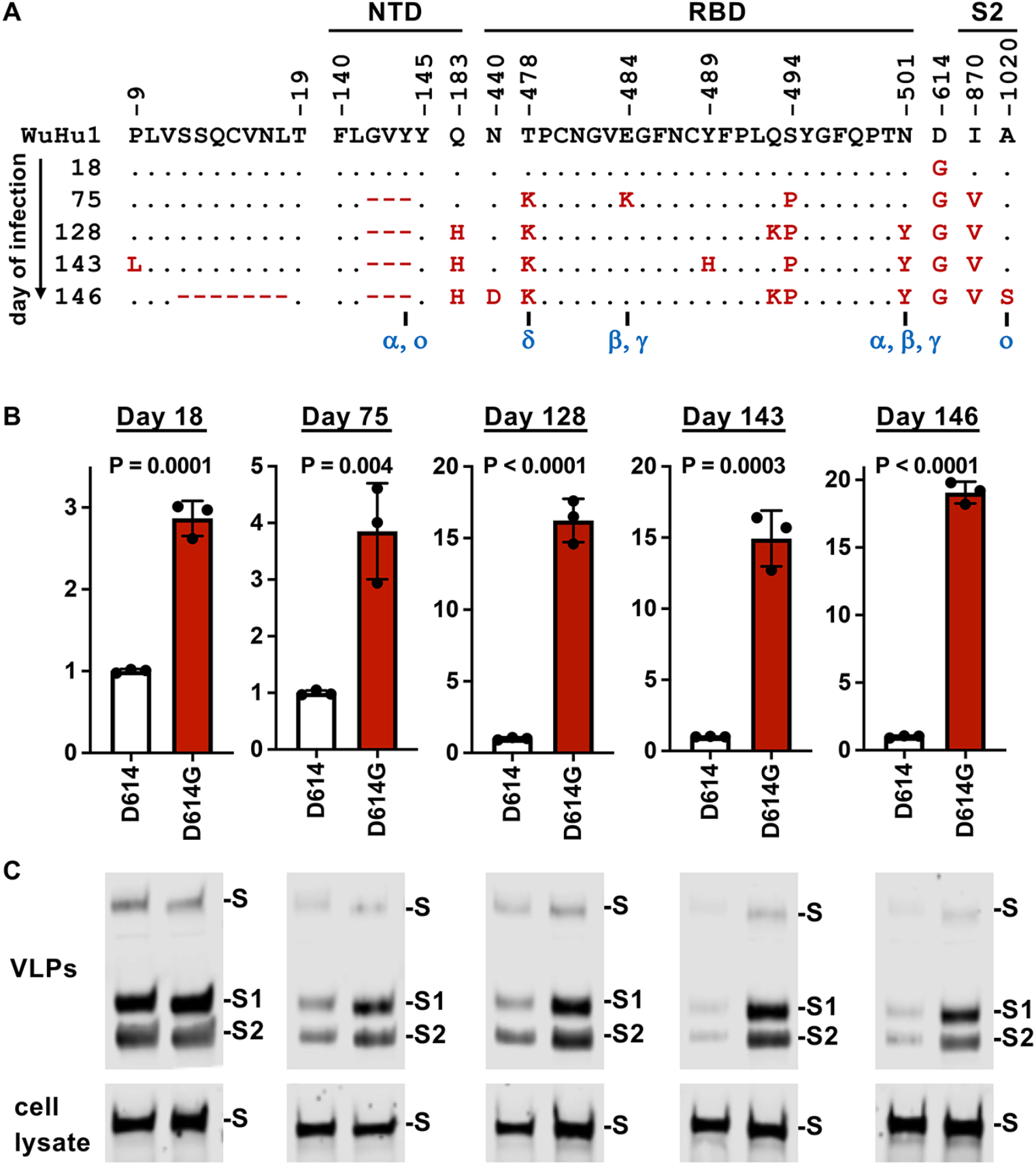
S:D614G-dependence of SARS-CoV-2 Spike variants isolated from an individual with prolonged infection. **(A)** Time-course of S mutations detected during prolonged SARS-CoV-2 infection in an immunosuppressed individual^48^. The day post-infection is indicated vertically on the left. Amino acids encoded at the earliest time-point (day 18) are indicated, as is their position within Spike. Aside from S:D614G, day 18 Spike residues were identical to those in Wuhan-Hu-1 (GenBank Accession NC_045512). Residues conserved at subsequent time-points are indicated with a dot, changed amino acids with red letters, and deletions with red dashes. Mutations observed in Variants of Concern are indicated with blue letters, using the WHO Greek letter designation. Corresponding Pango lineage names are B.1.1.7 (α), B.1.351 (β), P.1 (γ), B.1.617.2 (δ), and B.1.1.529/BA.1 (ο). **(B)** Infectivity of Spike mutations from the indicated days post-infection measured as pseudotypes with an HIV-1-based luciferase reporter vector, on ACE2^+^/TMPRSS2^+^/HEK293T target cells, as done previously^20^. Spikes from each time-point were tested with the actual S:D614G (red bars) and with the residue reverted to the ancestral S:D614 (white bars). Shown is one representative of three biological replicates, each of which used vectors produced by independent transfection. P values are two-tailed, unpaired T test in triplicate. **(C)** Steady-state levels of the indicated variant Spike proteins by western blot with a combination of antibodies specific for S1 and S2. Results are shown for the cell lysate (bottom panel) and for virus-like particles (VLPs) pelleted from the supernatant by ultracentrifugation (upper panel). Blots are representative of three independent transfections.

### SARS-CoV-2 S:D614G promotes the infectivity of Spikes from Variants of Concern

Two observations suggest that S:D614G was critical to the fitness of the complex combinations of S mutations that appeared in SARS-CoV-2 Variants of Concern. First, variants bearing large numbers of nonsynonymous S mutations were not detected until well after S:D614G had become prevalent worldwide^19, 20^. Second, at least four amino acid changes in the S:D614G-dependent SARS-CoV-2 Spike proteins isolated from an individual with prolonged infection were shared with Variants of Concern (Figure 1).

To determine whether the infectivity of Spike proteins encoded by Variants of Concern are S:D614G-dependent, plasmids were engineered to express Spike proteins of the Alpha (Pango lineage B.1.1.7), Beta (B.1.351), Gamma (P.1), Delta (B.1.617.2), and Epsilon (B.1.427) variants (Supplementary Figure 1). For each variant Spike, the relative infectivity of lentiviral vectors was assessed, comparing the wild-type S:D614G version with a mutant reverted to the ancestral S:D614. As previously reported for Spike on the Wuhan-Hu-1 background^20^, introduction of S:D614G increased infectivity 7.6-fold (Figure 2A). Mutation of S:D614G to the ancestral S:D614 compromised the infectivity of Spikes encoded by each of the Variants of Concern to a much greater extent (Figure 2A): for the Alpha Spike, the reduction was 71.0-fold (p<0.0001); for Beta, 17.3-fold (p<0.0001); for Gamma, 10.3-fold (p=0.008); for Delta, 10.0-fold (p=0.03); and for Epsilon, 23.6-fold (p<0.0001). These results indicate that the infectivity of virions bearing Spike proteins from these SARS-CoV-2 Variants of Concern is highly-dependent upon S:D614G.

Western blots with antibodies specific for the S1 and S2 proteins were used to assess the effect of S:D614G on the steady-state level of Spike protein for the variants. When lysate from cells expressing the Spike protein was examined, S:D614G had minimal effect on Spike protein from Wuhan-Hu-1 or any of the variants (Figure 2B). In contrast, dramatic reduction in virion-associated S1 protein was detected (Figure 2B). In some cases virion-associated S2 protein was also decreased (Figure 2B). These western blots suggest that S:D614G maintains the infectivity of virions bearing Spike protein from Variants of Concern by stabilizing virion-associated S1 protein and in some cases maintaining Spike incorporation into virions.

**Figure 2.**
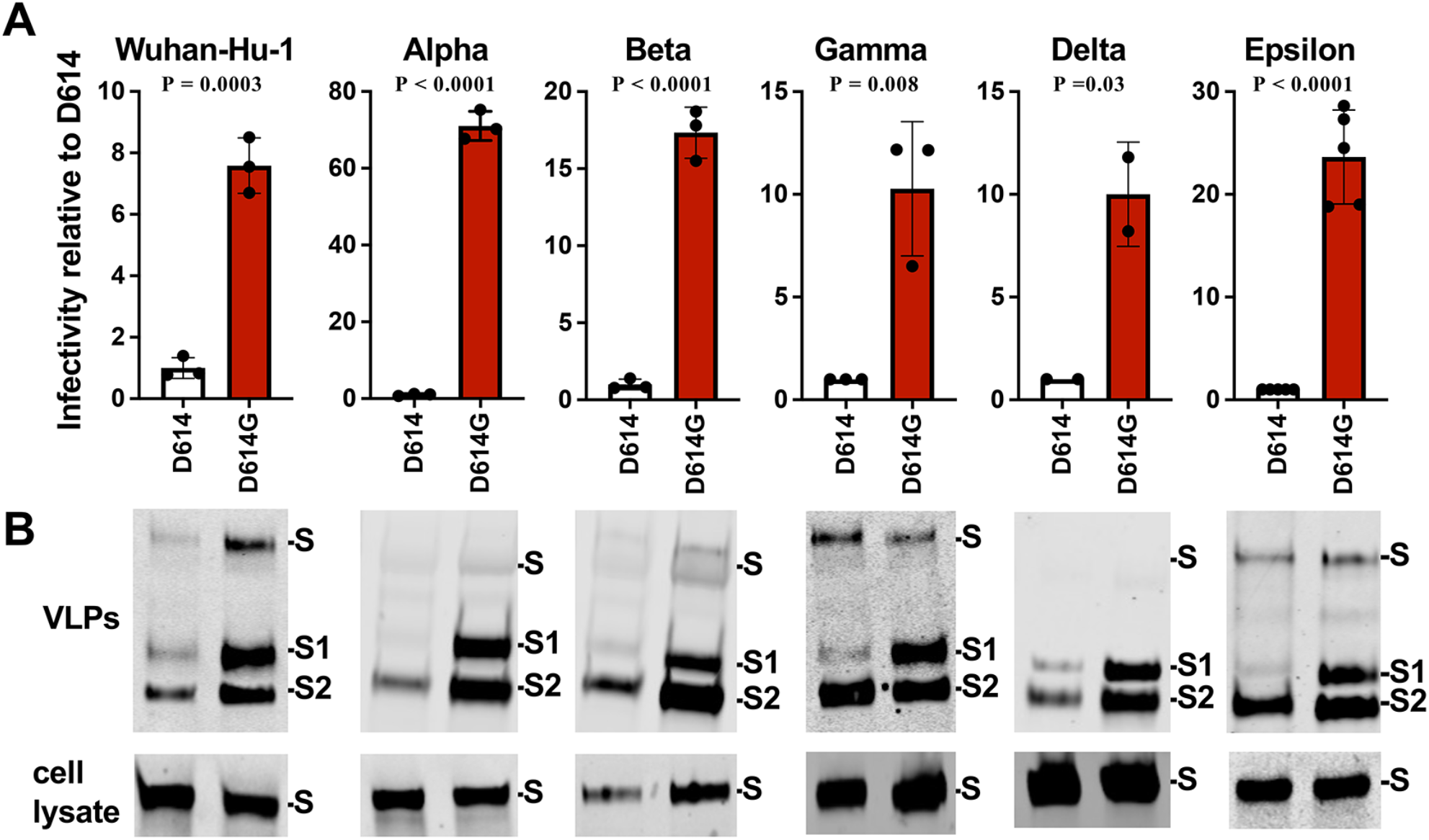
The infectivity of Spike proteins from Variants of Concern is highly-dependent upon S:D614G. **(A)** Infectivity of virions bearing Spike proteins from the indicated variants, assessed as pseudotypes with an HIV-1-based luciferase reporter vector, on ACE2^+^/TMPRSS2^+^/HEK293T target cells, as done previously^20^. Each variant of concern Spike was tested with its actual S:D614G (red bars) and with the residue reverted to the ancestral S:D614 present in Wuhan-Hu-1 (white bars). Data for each variant is reported as relative to S:D614 on the homologous background. Shown is one representative of three biological replicates, each of which used vectors produced by independent transfection. P values are two-tailed, unpaired T test in triplicate. **(B)** Steady-state levels of the indicated variant Spike proteins by western blot with a combination of antibodies specific for S1 and S2. Results are shown for the cell lysate (bottom panel) and for virus-like particles (VLPs) pelleted from the supernatant by ultracentrifugation (upper panel). Blots are representative of three independent transfections.

**Supplementary Figure 1.**
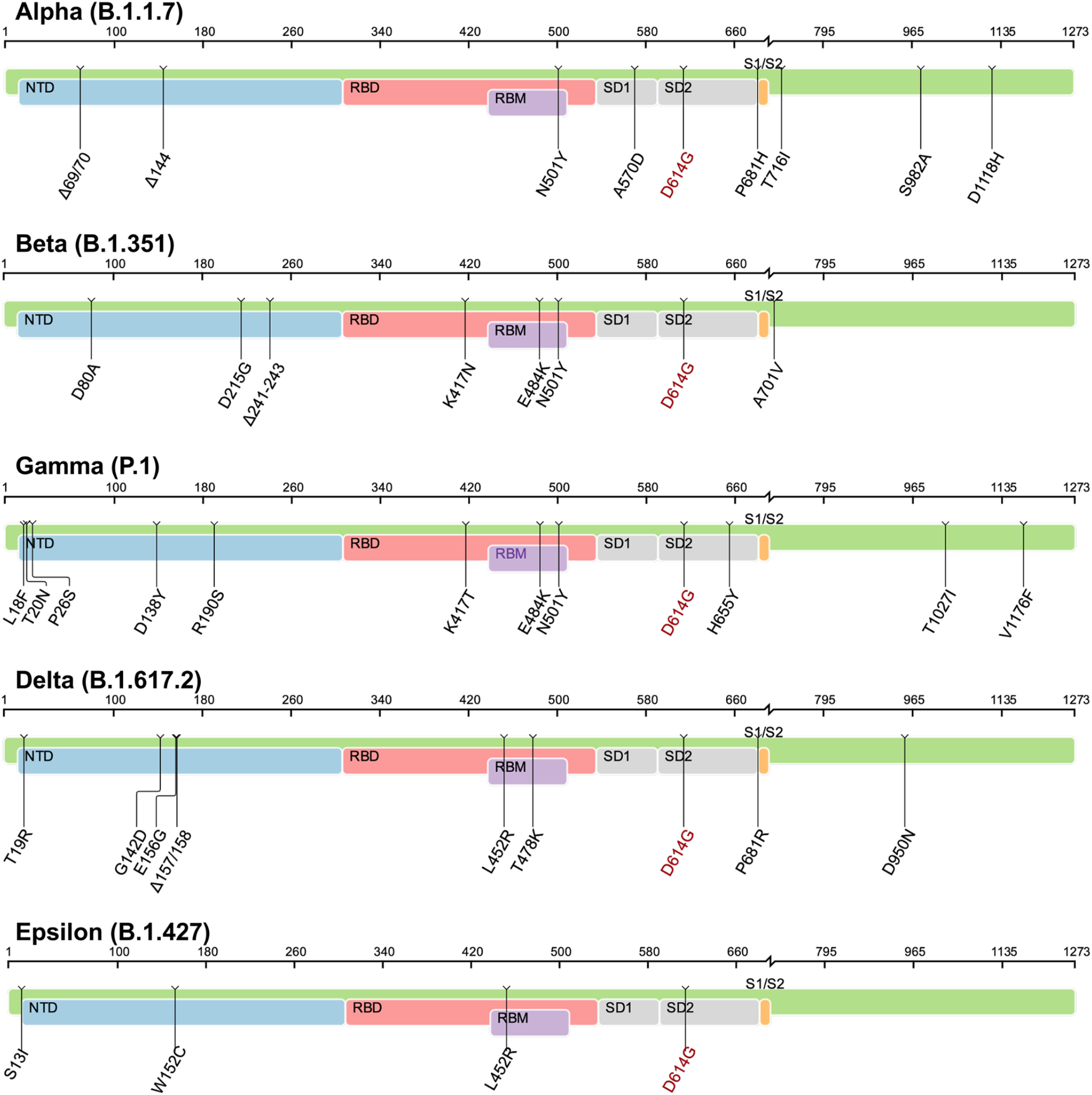
Schematic diagrams showing amino acid changes along the length of the indicated SARS-CoV-2 variant Spike proteins (related to Figure 2). Variant names are according to the WHO greek letter system. Pango lineage is in parentheses^62^. Amino acid numbering and amino acid changes are relative to the ancestral SARS-CoV-2 Wuhan-Hu-1 (GenBank Accession NC_045512). The S:D614G mutation that is common to all the variants is highlighted in red font. Diagrams provided by the Coronavirus Resistance Database^63^.

### SARS-CoV-2 variant infectivity in the absence of S:D614G

S:D614G is the single most common Spike mutation^33^ but some variants lacking S:D614G dominated regional outbreaks for more than one year after the start of the SARS-CoV-2 pandemic. The A.23.1 variant retained the ancestral S:D614 residue (Supplementary Figure 2) and accounted for 90% of the SARS-CoV-2 isolates in Uganda at the beginning of 2021 despite competition from multiple S:D614G-bearing B lineages^64^. This suggested that A.23.1 Spike possesses an amino acid which confers replication advantage similar to that of S:D614G. Reversion of S:Q613H in A.23.1 to the ancestral residue decreased infectivity 9.5-fold (Figure 3A). Thus, of the six Spike non-synonymous mutations that distinguish A.23.1 from Wuhan-Hu-1 (GenBank Accession NC_045512), S:Q613H is sufficient to promote infectivity to a similar extent as S:D614G. Western blot comparing the Spike signal for A23.1 with that of the reversion mutation showed that S:Q613H increased steady-state S1 protein levels on virions (Figure 3B), similar to the effect of S:D614G (Figure 2).

Another variant lacking S:D614G, A.27 (Supplementary Figure 2), circulated in 31 countries until June 2021^65^. Among the seven non-synonymous S mutations in A.27, S:L18F, S:L452R, S:N501Y, and S:H655Y are shared with more than one Variant of Concern. Reversion of S:H655Y to the ancestral residue caused a 10.7-fold decrease in infectivity (Figure 3C), as well as a profound decrease in steady-state S1 and S2 proteins associated with virions (Figure 3D), indicating that this mutation is sufficient to maintain the infectivity of A.27.

The A.30 variant (Supplementary Figure 2) was discovered in a screen of travelers arriving at an Angolan airport a year before the appearance of Omicron^42^. It lacked S:D614G but possessed 16 nonsynonymous S mutations. Like A.27, the infectivity of A.30 was dependent upon S:H655Y. Reversion of S:H655Y to the ancestral residue caused an 11-fold decrease in infectivity (Figure 3E), and disrupted steady-state levels of S1 and S2 proteins associated with virions (Figure 3F).

The effect of S:H655Y was then compared with that of S:D614G on different Spike backgrounds. On the original Wuhan-Hu-1 Spike, the S:D614G mutation caused a 6.4-fold increase in infectivity, while S:H655Y caused a 7.7-fold increase in infectivity (Figure 3G). When the two mutations were combined there was a 15.5-fold increase in infectivity (Figure 3G). Western blot showed a modest increase in virion-associated S protein in the presence of either mutation (Figure 3H). On the Wuhan-Hu-1 Spike background, then, S:D614G and S:H655Y both increased infectivity to a similar extent, and the effect of the two mutations was additive.

The Gamma Variant possessed both the S:D614G and S:H655Y mutations (Supplementary Figure 1). Each of these residues was reverted to the ancestral version and the effect of each amino acid change was assessed. S:D614G alone caused a 21.9-fold increase in infectivity on the Gamma background, whereas S:H655Y caused a 3.9-fold increase (Figure 3I). When the two mutations were combined the infectivity increased 36.9-fold (Figure 3I). In the case of the Gamma Variant, S1 protein levels on the virion were dependent upon either mutation though the effect of S:D614G was larger (Figure 3J). The infectivity of the Gamma Variant was more dependent on S:D614G than S:H655Y, though the effect of the two mutations together was still significantly greater than the effect of S:D614G alone (Figure 3I, p = 0004).

Like the Gamma Variant, the Omicron BA.1 Variant also possessed both the S:D614G and S:H655Y mutations (Supplementary Figure 2). On the BA.1 background. S:D614G increased infectivity 5.1-fold and S:H655Y increased infectivity 48.1-fold (Figure 3K). When the two mutations were combined together the infectivity increased 51.3-fold (Figure 3K). In the case of BA.1, S:D614G and S:H655Y both increased S1 protein levels on the virion (Figure 3L). As compared with Wuhan-Hu-1 and the Gamma Variant, BA.1 was more dependent upon both mutations for full infectivity. In contrast to the Gamma Variant, the infectivity of the Omicron (BA.1) variant was more dependent on S:H655Y than on S:D614G.

**Figure 3.**
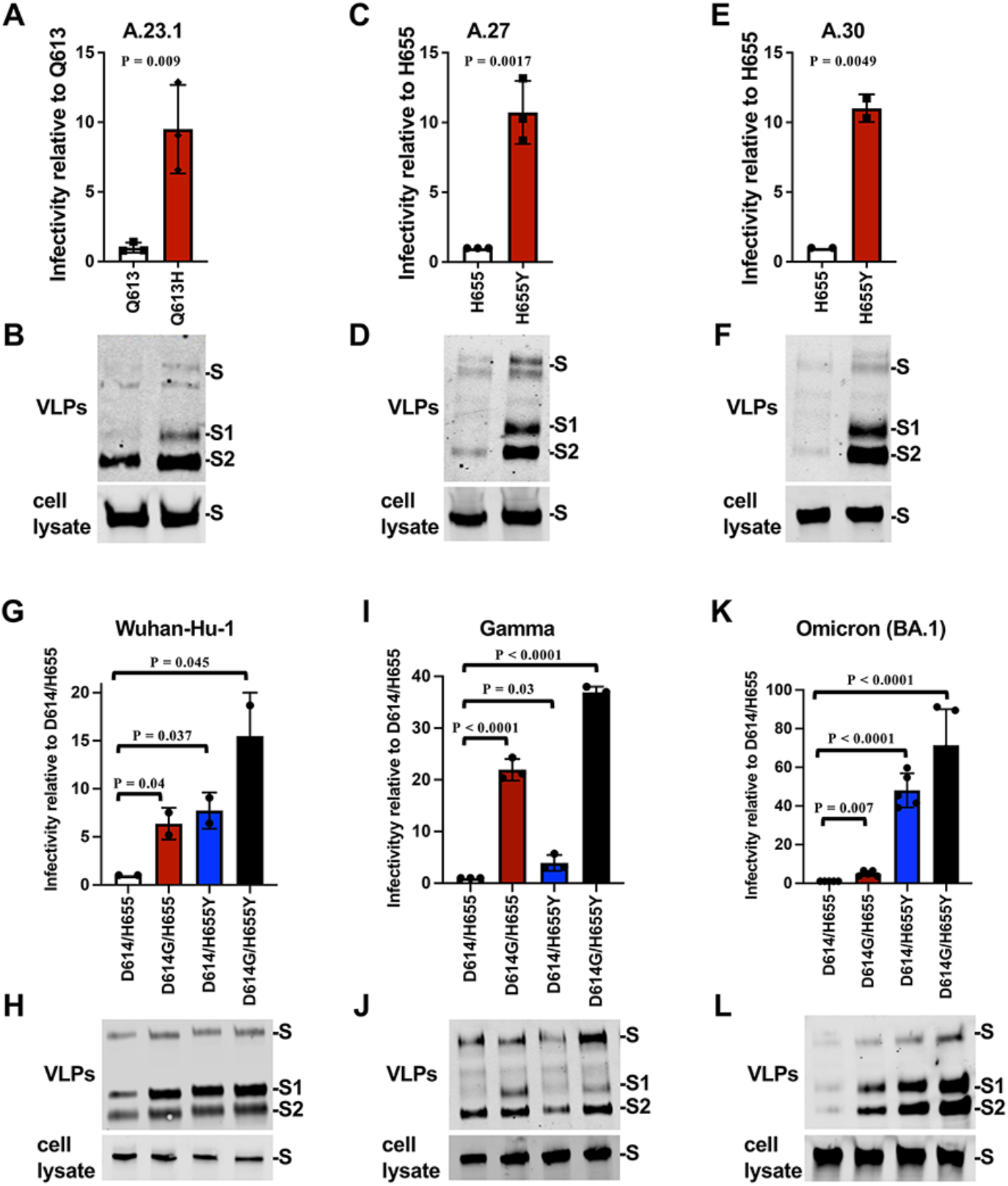
Convergent evolution of SARS-CoV-2 mutants S:D614G, S:Q613H, and S:H655Y. **(A)** Infectivity of virions bearing A.23.1 Spike protein, either wild-type (S:Q613H) or the S:Q613 reversion mutation, assessed as pseudotypes on an HIV-1-based luciferase reporter vector, on ACE2^+^/TMPRSS2^+^/HEK293T target cells, as done previously ^20^. Data is reported as relative to S:Q613. P values are for two-tailed, unpaired T test in triplicate. **(B)** Steady-state levels of the indicated variant Spike proteins by western blot with a combination of antibodies specific for S1 and S2. Results are shown for the cell lysate (bottom panel) and for virus-like particles (VLPs) pelleted from the supernatant by ultracentrifugation (upper panel). Blots are representative of three independent transfections. **(C)** Infectivity of virions pseudotyped with A.27 Spike, either wild-type (S:H655Y) or the S:H655 reversion mutation, assessed as in (A). **(D)** Western blot of A.27 Spike, either wild-type (S:H655Y) or the S:H655 reversion mutation, assessed as in (B). **(E)** Infectivity of virions pseudotyped with A.30 Spike, either wild-type (S:H655Y) or the S:H655 reversion mutation, assessed as in (A). **(F)** Western blot of A.30 Spike protein, either wild-type (S:H655Y) or the S:H655 reversion mutation, assessed as in (B). **(G)** Infectivity of virions pseudotyped with the Wuhan-Hu-1 Spike bearing the indicated mutations, assessed as in (A). **(H)** Western blot of Wuhan-Hu-1 Spike protein bearing the indicated mutations, assessed as in (B). **(I)** Infectivity of virions pseudotyped with the Gamma (P.1) Spike bearing the indicated mutations, assessed as in (A). **(J)** Western blot of Gamma (P.1) Spike bearing the indicated mutations, assessed as in (B). **(K)** Infectivity of virions pseudotyped with the Omicron (BA.1) Spike bearing the indicated mutations, assessed as in (A). **(L)** Western blot of the Omicron (BA.1) Spike protein bearing the indicated mutations, assessed as in (B).

**Supplementary Figure 2.**
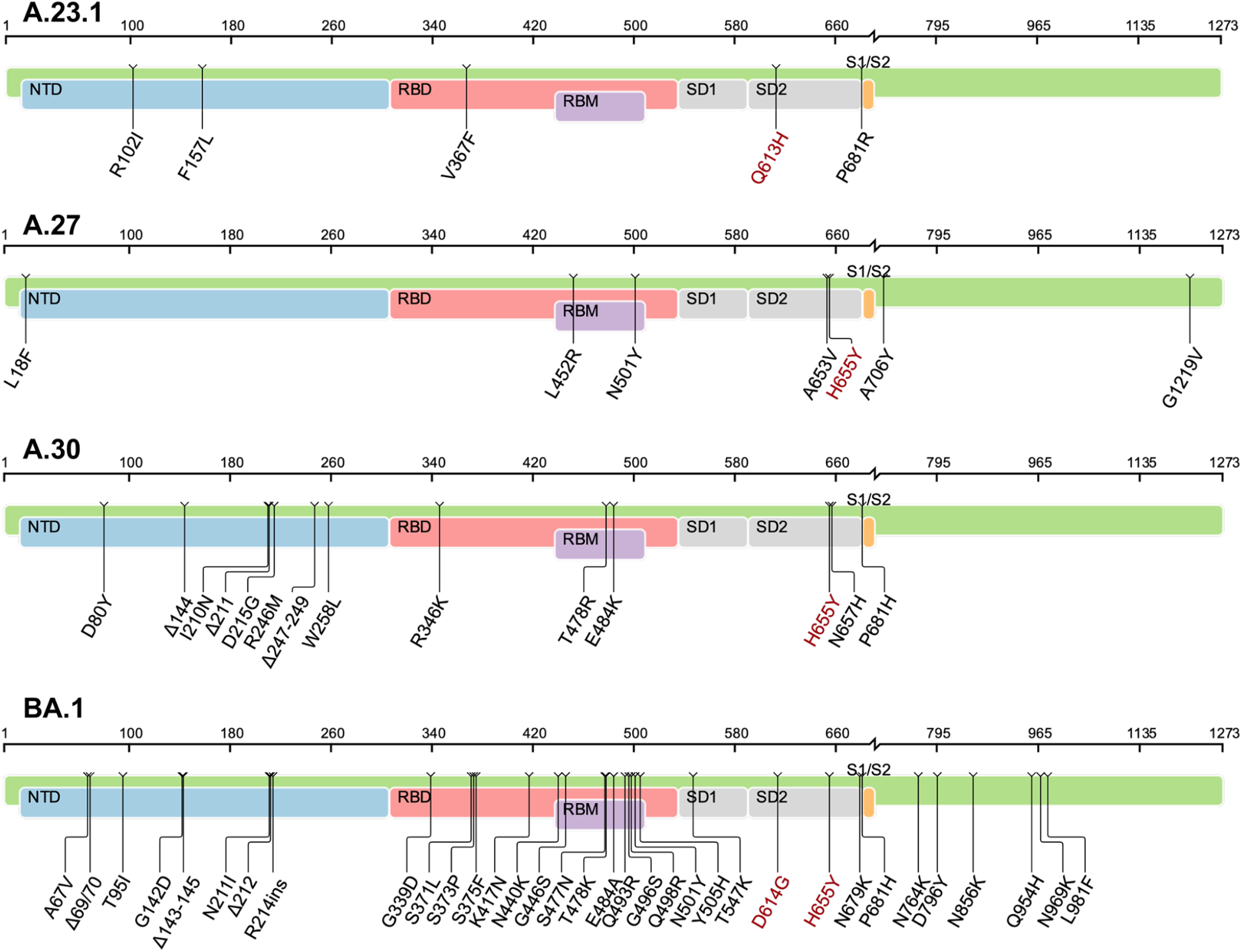
Schematic diagrams showing amino acid changes along the length of the indicated SARS-CoV-2 variant Spike proteins (related to Figure 3). Amino acid numbering and amino acid changes are relative to the ancestral SARS-CoV-2 Wuhan-Hu-1 (GenBank Accession NC_045512). Diagrams provided by the Coronavirus Resistance Database^63^.

### Infectivity of authentic SARS-CoV-2 Omicron (BA.1) is dependent on both S:D614G and S:H655Y

The contribution of S:D614G and S:H655Y to the infectivity of authentic Omicron BA.1 virus was evaluated next. Recombinant SARS-CoV-2 Omicron (BA.1) was engineered to encode the ancestral Wuhan-Hu-1 amino acid residues at positions 614 and 655, individually and in combination^66, 67^. The plaque size on TMPRSS2^+^ Vero-E6 cells of Omicron (BA.1) wildtype (S:D614G/H655Y), S:D614/H655Y, S:D614G/H655, and S:D614/H655 was comparable (Figure 4A). Specific infectivity of the first passage, sequence-verified stocks of each recombinant virus was determined by calculating the ratio of SARS-CoV-2 RNA to plaque-forming units (PFU). As compared to the wild-type (S:D614G/H655Y), the infectivity of S:D614/H655Y was reduced 9.4-fold, S:D614G/H655 was reduced 13.8-fold, and the double revertant S:D614/H655 was reduced 172.4-fold (Figure 4B). Spike proteins encoded by the four recombinant viruses were assessed by western blot after infecting TMPRSS2^+^ Vero-E6 cells for 48 hrs with an equal multiplicity of infection for each recombinant. At the end of the collection period, the identity of each virus was confirmed by next-generation sequencing. The steady-state levels of Spike protein in cell lysate, the degree of proteolytic processing, and the amount of Spike associated with cell-free virions decreased in accordance with the decrease in infectivity (Figure 4C).

The ability of wild-type (S:D614G/H655Y) SARS-CoV-2 Omicron (BA.1) to compete against each of the recombinant viruses bearing reversion mutations was assessed next using primary human tracheal and bronchial epithelial cells in an air-liquid interface model (HAE) that has been used previously to compare the replication SARS-CoV-2 variants^68, 69^. The wild-type Omicron BA.1 was mixed with a 3:1 ratio of each mutant based on PFU/ml and used to infect the culture for two hours. Cells were washed and media was removed from the apical surface at the airway interface. Each day for three days, virus was collected by bathing the apical surface in media and the ratio of wild-type to mutant virus was determined by next-generation sequencing. Despite greater abundance of mutant virus RNA at time zero for each of the three challenges, by the end of three days the wild-type virus dominated each of the cultures (Figure 4D). S:D614G/H655 was more efficiently outcompeted by the wild-type than was S:D614/H655Y, and the double reversion mutant S:D614/H655 was more efficiently outcompeted than either of the single mutants (Figure 4D). In agreement with the single cycle infectivity data (Figure 3K), these data demonstrate that S:D614G and S:H655Y both contribute to the infectivity of authentic, replication-competent Omicron SARS-CoV-2 (BA.1).

**Figure 4.**
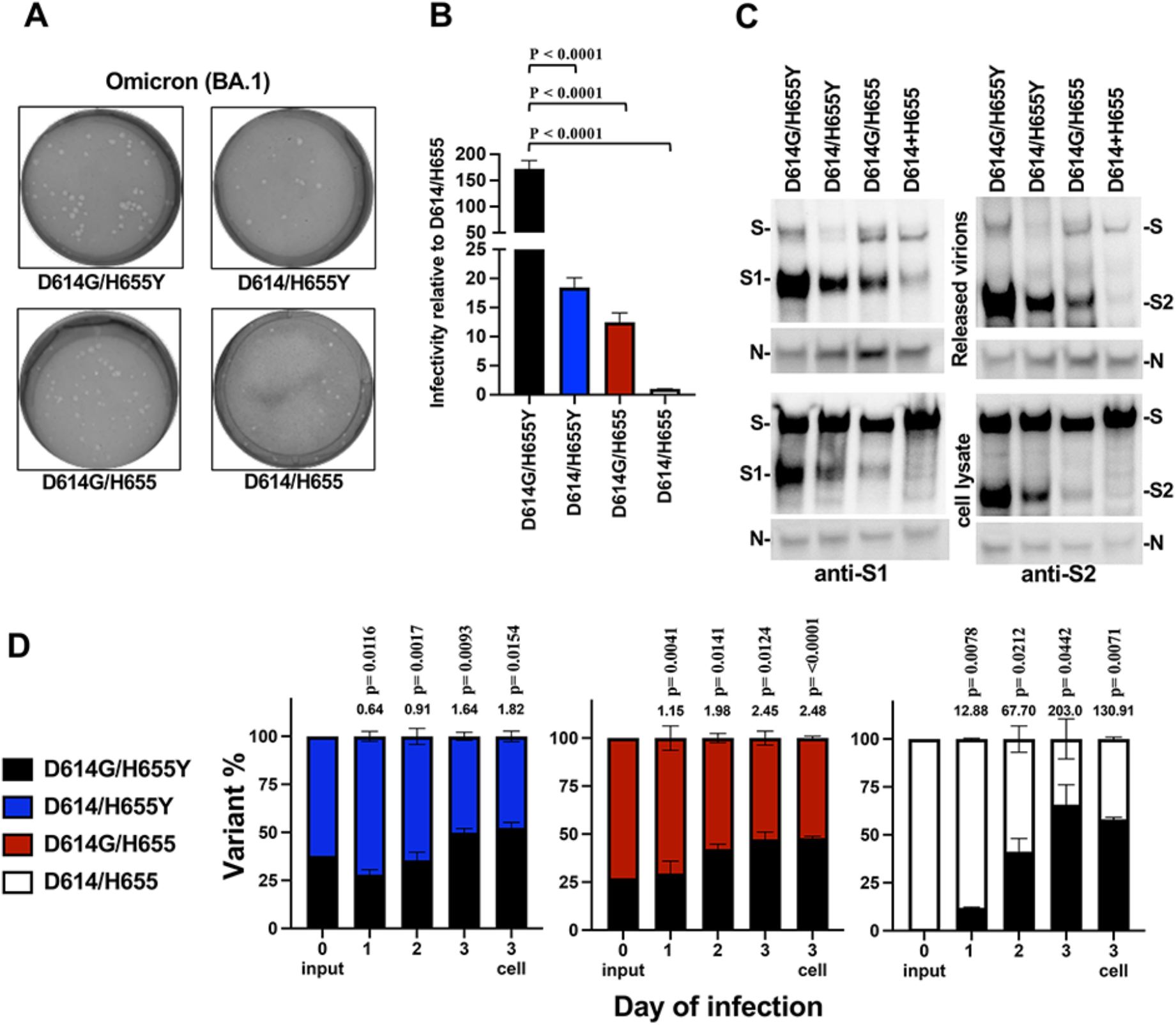
S:D614G and S:H655Y both contribute to infectivity of authentic Omicron (BA.1) SARS-CoV-2. **(A)** Plaque morphology of recombinant Omicron (BA.1), either wild type (S:D614G/H655Y), S:D614/H655Y, S:D614G/H655, or S:D614/H655, on Vero E6 cells expressing TMPRSS2. (**B**) Specific infectivity of P1 stocks for each recombinant virus, relative to S:D614/H655, as determined by the ratio of SARS-CoV-2 RNA by RT-qPCR to PFU. The graphs show the mean with 95% confidence interval of 4 technical replicates each for 4 biological replicates. P values were determined by one-way ANOVA with Dunnett’s multiple comparisons test. (**C**) Western blot of the indicated SARS-CoV-2 Omicron (BA.1) recombinants. TMPRSS2-expressing Vero E6 cells were infected with 0.1 MOI of the indicated recombinant. Producer cell lysate and cell-free virions were probed with antibodies against S1 (left) and S2 (right). Loading of cell lysate was normalized with anti-N antibody. A representative of three experiments is shown. (**D**) Head-to-head competition between Omicron (BA.1) recombinants. For each competition, wild type (S:D614G/H655Y) was mixed with one of the indicated mutants in a 3:1 ratio according to PFU calculations. The mixture was then used to challenge primary human tracheal/bronchial epithelial cells in an airway 3D mucociliary tissue model (HAE) at an MOI of 0.2. Secreted progeny viruses were collected daily from the apical side of the culture and on day 3 cellular RNA was collected. The percentage of each variant in input and days 1, 2 and 3 post-infection was determined by NGS. For each time point, data is represented as percent (%) of each variant present. The number above each bar represents the relative fitness as normalized to the input ratio. P values are calculated as the coefficient of each linear regression analysis of RNA ratio at a given time-point versus input RNA ratio.

### SARS-CoV-2 S:D614G and S:H655Y are adaptations to the polybasic S1/S2 cleavage site

Rapid increase in S:D614G prevalence over the first six months of the SARS-CoV-2 pandemic suggests that this mutation was an adaptation selected by person-to-person transmission after spillover from an animal host. S:D614G appears not to be an adaptation to the human ACE2 ortholog since it increased SARS-CoV-2 infectivity to a similar extent on cells bearing ACE2 orthologs from horseshoe bats, pangolins, and other potential intermediate hosts^20^.

It is puzzling that S:D614G has only been detected once among other members of the *Sarbecovirus* subgenus (GenBank accession #FJ882963), which includes the zoonotic human pathogen SARS-CoV-1. Like SARS-CoV-2, SARS-CoV-1 enters cells via ACE2^70^. SARS-CoV-2 is distinguished from other Sarbecoviruses by the presence of a polybasic amino acid cleavage site at the Spike S1/S2 junction. This raises the possibility that S:D614G was selected during human-to-human transmission as an adaptation to this molecular feature. Consistent with this hypothesis, S:D614G caused no significant increase in SARS-CoV-1 Spike infectivity unless it was also engineered to include the SARS-CoV-2 polybasic S1/S2 cleavage site (Figure 5A and B). Conversely, disruption of the S1/S2 cleavage site of the Wuhan-Hu-1 or Alpha Variant Spikes, rendered infectivity of these Spikes independent of S:D614G (Figure 5C-F), as did an inhibitor of proprotein convertases (Figure 5G and H).

Additional evidence that the polybasic S1/S2 cleavage site drove the observed evolution of the SARS-CoV-2 Spike was sought by modifying the Wuhan-Hu-1 Spike to include the S:P681R mutation. The latter mutation was present in the Delta Variant, which also had S:D614G, and in A.23.1, which had S:Q613H (Supplementary Figure 1 and 2). Similar increase in cleavage efficiency was seen with S:P681H, a mutation present in the Alpha Variant, which had S:D614G, in the Omicron BA.1 Variant, which had both S:D614G and S:H655Y, and in A.30, which had S:H655Y (Supplementary Figures 1 and 2). In the presence of S:P681R, the S:Q613H, S:D614G, and S:H655Y mutations increased the infectivity of the Wuhan-Hu-1 Spike 4-fold, 31.8-fold, and 40.3-fold, respectively (Figure 5I and J). This magnitude increase was much greater than was observed in the absence of S:P681R (for example, as compared with Figure 3G and H), indicating that, in the presence of a more efficient S1/S2 cleavage site, these mutations have a greater effect on infectivity.

While the polybasic S1/S2 cleavage site is dispensable and even selected against during propagation on commonly used cell lines^25, 71^, it is required for entry into human lung and primary airway epithelial cells, transmission in ferrets, for pathogenesis in small animal models, and rarely is it deleted during human-to-human transmission ^72–74^. When recombinant Omicron (BA.1) was replicated on TMPRSS2^+^ Vero-E6 cells for 48 hrs the sequence of the virus was identical to the input virus (Figure 4). In contrast, when target cells were TMPRSS2-negative Vero-E6 cells, the Omicon (BA.1) S:D614/H655 double revertant mutant virus acquired a 24-nucleotide deletion mutation that removed the polybasic S1/S2 cleavage site (Figure 5K); in side-by-side experiments the S1/S2 cleavage site remained intact in the wild type (S:D614G/H655Y), S:D614/H655Y, and S:D614G/H655 recombinants. When this polybasic S1/S2 cleavage site mutation was combined with the S:D614/H655 double mutant in the lentiviral pseudotype assay it rescued single-cycle infectivity to the level of the wild-type Omicron (BA.1) S:D614G/H655Y (Figure 5L). Taken together with the above results, these data provide evidence that D614G and H655Y are adaptations to the polybasic S1/S2 cleavage site.

**Figure 5.**
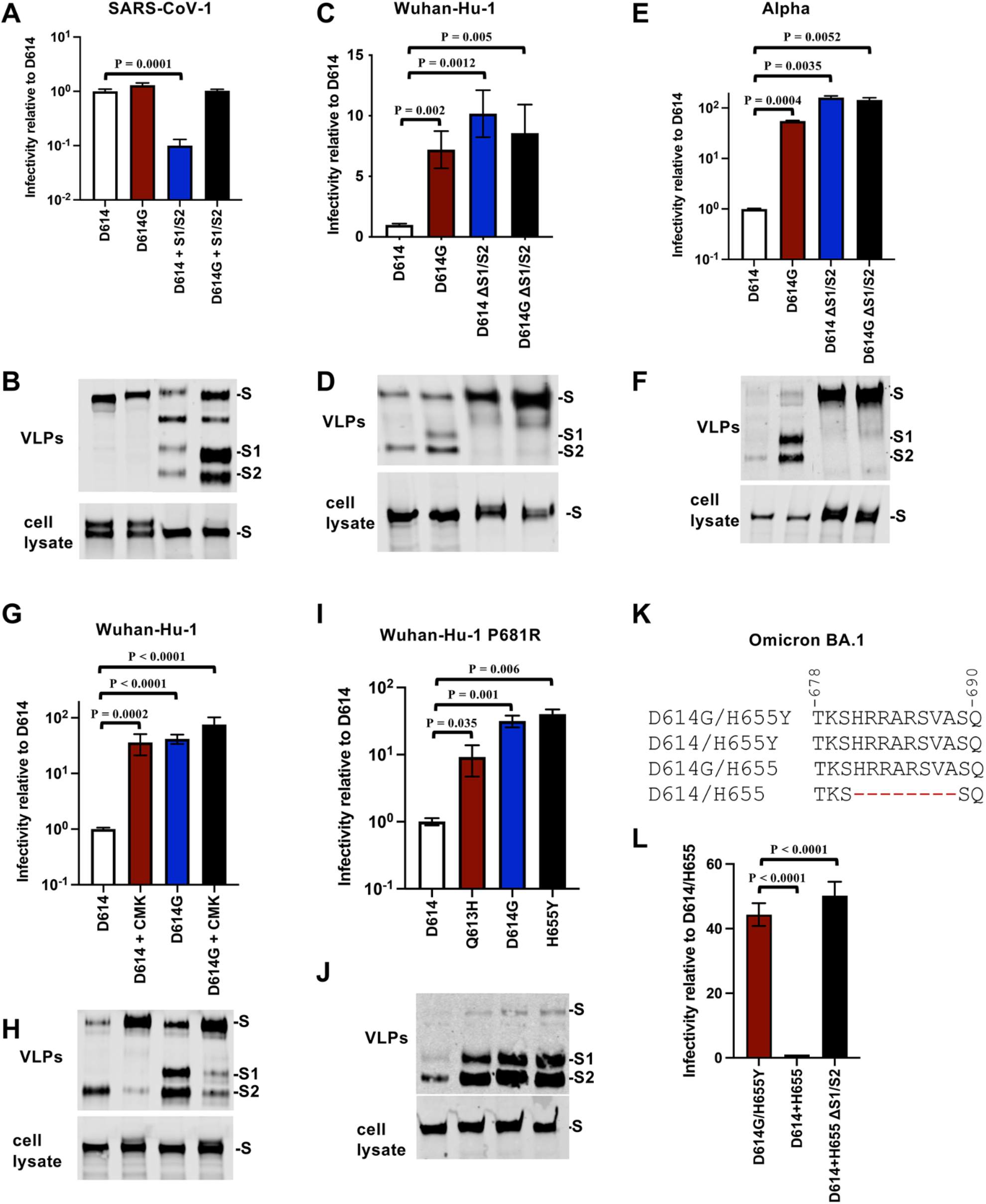
S:D614G and S:H655Y phenotype dependence on the polybasic S1/S2 cleavage site. **(A)** Infectivity of virions bearing the SARS-CoV-1 Spike protein, either wild-type (S:D614) or S:D614G, with or without the S1/S2 cleavage site from SARS-CoV-2, assessed as pseudotypes with an HIV-1-based luciferase reporter vector, on ACE2^+^/TMPRSS2^+^/HEK293T target cells, as done previously^20^. Data is reported as relative to S:D614. P values are for two-tailed, unpaired T test in triplicate. **(B)** Steady-state levels of the indicated variant Spike proteins by western blot with a combination of antibodies specific for S1 and S2. Results are shown for the cell lysate (bottom panel) and for virus-like particles (VLPs) pelleted from the supernatant by ultracentrifugation (upper panel). Blots are representative of three independent transfections. **(C)** Infectivity of virions pseudotyped with SARS-CoV-2 Wuhan-Hu-1 Spike, either wild-type (S:D614) or S:D614G, with or without disruption of the S1/S2 cleavage site, assessed as in (A). **(D)** Western blot of SARS-CoV-2 Wuhan-Hu-1 Spike, either wild-type (S:D614) or S:D614G, with or without disruption of the S1/S2 cleavage site, assessed as in (B). **(E)** Infectivity of virions pseudotyped with the SARS-CoV-2 Alpha Variant Spike, either wild-type (S:D614G) or S:D614, with or without disruption of the S1/S2 cleavage site, assessed as in (A). **(F)** Western blot of SARS-CoV-2 Alpha Variant Spike, either wild-type (S:D614G) or S:D614, with or without disruption of the S1/S2 cleavage site, assessed as in (B). **(G)** Infectivity of virions pseudotyped with SARS-CoV-2 Wuhan-Hu-1 Spike, either wild-type (S:D614) or S:D614G, with or without the proprotein convertase inhibitor CMK, assessed as in (A). **(H)** Western blot of SARS-CoV-2 Wuhan-Hu-1 Spike, either wild-type (S:D614) or S:D614G, with or without the proprotein convertase inhibitor CMK, assessed as in (B). **(I)** Infectivity of virions pseudotyped with the SARS-CoV-2 Wuhan-Hu-1 Spike bearing S:P681R, and either S:D614, S:Q613H, S:D614G, or S:H655Y, assessed as in (A). **(J)** Western blot of SARS-CoV-2 Wuhan-Hu-1 Spike bearing S:P681R, and either S:D614, S:Q613H, S:D614G, or S:H655Y, assessed as in (B). **(K)** Amino acid sequence at the S1/S2 cleavage site encoded by recombinant Omicron (BA.1) bearing either S:D614G/H655Y (wild-type), S:D614/H655Y, S:D614G/H655, or S:D614/H655, after propagation of P1 stocks on Vero-E6 cells for 48 hrs. **(L)** Infectivity of virions pseudotyped with the SARS-CoV-2 Omicron (BA.1) Spike, either S:D614G/H655Y (wild-type), S:D614/H655, or S:D614/H655 in combination with the 8 amino acid deletion flanking the S1/S2 cleavage site shown in (K), assessed as in (A).

**Supplementary Figure 3.**
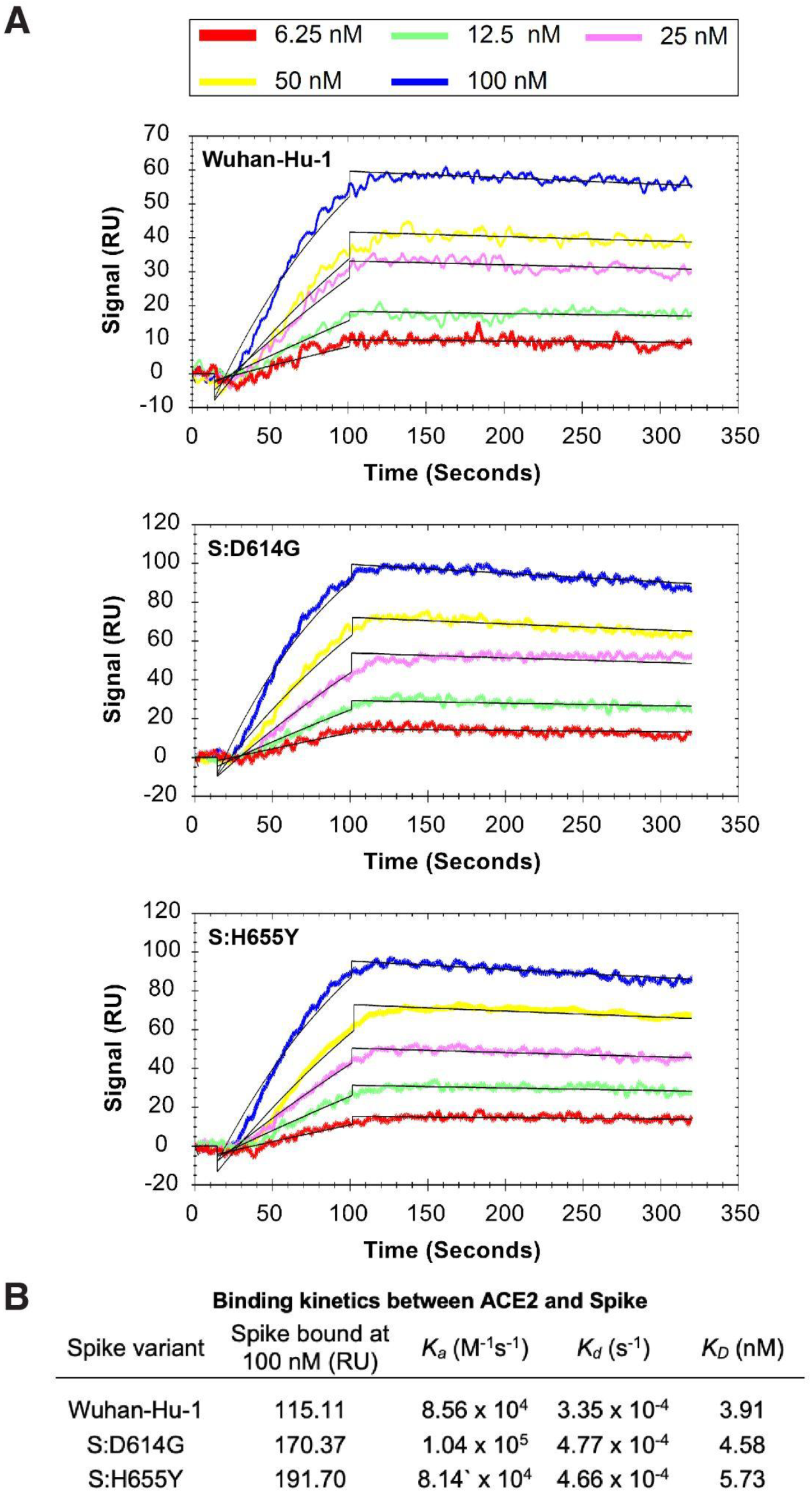
SARS-CoV-2 Spike variant binding kinetics to ACE2. (**A**) Spike variants binding to ACE2 measured by SPR. Kinetic parameters were obtained by fitting the data to a 1:1 binding model (black lines). **(B)** Summary of kinetic parameters measured in (A).

### Like S:D614G, S:H655Y shifts the receptor binding domain of Wuhan-Hu-1 Spike to an open conformation

S:D614G and S:H655Y might increase SARS-CoV-2 infectivity by strengthening ACE2 affinity. Surface plasmon resonance was used here to assess ACE2 binding kinetics for the Wuhan-Hu-1, S:D614G, and S:H655Y spikes (Supplementary Figure 3). Soluble, intact, homotrimeric preparations of the three Spike proteins were produced in human cells, and enriched using Ni-NTA resin and size-exclusion chromatography (Supplementary Figure 4A and 4B). In agreement with previous reports^20, 75^, Spike bearing S:D614G bound less tightly to ACE2 than did Wuhan-Hu-1 Spike (S:D614 *K_D_* = 3.91 nM; S:D614G *K_D_* = 4.58 nM), due to increase in the dissociation rate (S:D614 *k_d_* = 3.35 x 10^-4^ s^-1^; S:D614G *k_d_* = 4.77 x 10^-4^ s^-1^). Similar decrease in ACE2 binding affinity, and increase in dissociation rate, was observed with S:H655Y (*K_D_* = 5.73 nM; *k_d_* = 4.66 x 10^-4^ s^-1^) (Supplementary Figure 3). These results indicate that S:D614G and S:H655Y each bind less tightly to ACE2 than does the Wuhan-Hu-1 Spike, due to a faster off-rate.

The receptor-binding domain (RBD) on each protomer of the SARS-CoV-2 Spike trimer can adopt either a closed or open conformation^76, 77^. The ACE2 binding interface is concealed when the RBD is in the closed conformation^78, 79^ and only becomes accessible for ACE2-binding upon transition to the open conformation^80^. Previous studies showed that S:D614G increases the probability that the RBDs in the Wuhan-Hu-1 Spike trimer are in the open conformation^20^. The effect of S:H655Y on the SARS-CoV-2 Wuhan-Hu-1 Spike protein conformation was therefore examined using cryo-electron microscopy (cryo-EM).

The Spike proteins produced above were subjected to a previously established cryo-EM workflow (Supplementary Figure 3C). Well-defined particles were identified (Supplementary Figure 3D), and following particle extraction and reference-free two-dimensional classification, structural details of the S protein homotrimer were observed (Supplementary Figure 3E). Three-dimensional clustering and refinement revealed a single density map for S:H655Y (Figure 6A), with general architecture similar to previously published maps. Fourier shell correlation (FSC) analysis indicated that the map had a mean resolution of 4.1 Å (gold-standard criteria; Supplementary Figure 3F). Resolution in the area of the map where S:H655Y is located ranged from 3.5 to 4.0 Å (Supplementary Figure 3H).

A structural model was built using a previously deposited SARS-CoV-2 S protein model (PDB: 6VYB) as a starting point, and the model was validated based on the cryo-EM density map (Figure 6A, Supplementary Figure 3G, and Supplementary Table 1). Clustering analysis revealed that 100% of S:H655Y trimers had one RBD in the open conformation (Figure 6B). In contrast, in wild-type Wuhan-Hu-1 Spike, only 47% of Spike trimers had one RBD in the open conformation and 53% of trimers had all three RBDs in the closed state (Figure 6B). This shift of S:H655Y towards a more open RBD conformation, on-pathway for ACE2 binding, is reminiscent of the effect of S:D614G^20^ (Figure 6B).

In wild-type Wuhan-Hu-1 Spike, S:H655 points away from the protein towards the solvent (Figure 6C). In contrast, S:H655Y flips inward to nestle within a pocket formed by residues from the C-terminal domain of S1 (F643, N657, and I670) and from S2 (A694 and T696) (Figure 6C). The distances between amino acid 655 and these neighboring residues were shorter in S:H655Y than in the Wuhan-Hu-1 Spike (Figure 6D). Importantly, some of these distances are shorter in both the closed and open protomers in the complex, indicating that these residues maintain a stably-packed CTD regardless of RBD conformation. Accordingly, the tighter packing of S:H655Y may facilitate polar and hydrophobic interactions with residues that would not be possible with the Wuhan Hu-1 Spike (Figure 6D). Interestingly, one of the interactions introduced by H655Y is a polar interaction with T696. This interaction bridges the S1 and S2 polypeptide chains (H655Y on S1; T696 on S2) and would stabilize the CTD, making it less prone to dissociation following proteolytic processing at the S1/S2 cleavage site. Importantly, these interactions exist in both the closed and open protomers in the complex, indicating that these residues maintain a stably-packed CTD regardless of the conformation of the RBD.

**Figure 6.**
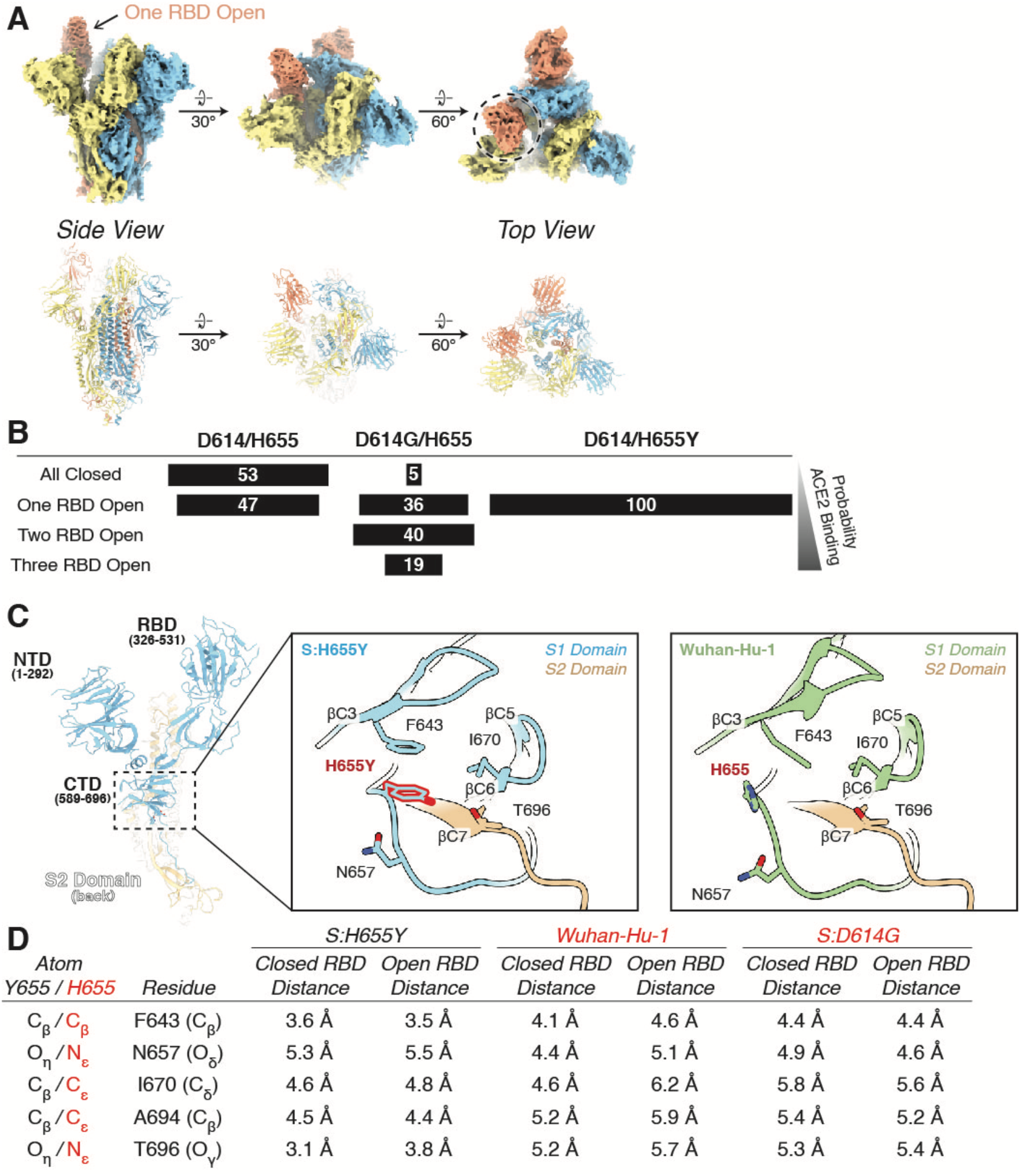
Cryo-EM structural determination of S:H655Y. **(A)** Cryo-EM density map (EMD-29910) and atomic model (PDB: 8GB0) for the S:H655Y trimer. **(B)** Summary of distributions of SARS-CoV-2 Wuhan-Hu-1, S:D614G, and S:H655Y conformations observed by cryo-EM. **(C)** Y655 flips inwards and interacts with other residues located within the C-terminal Domain (CTD). **(D)** Summary of distances between Y655/H655 and other residues within the CTD for S:H655Y, Wuhan-Hu-1, and S:D614G.

**Supplementary Figure 4.**
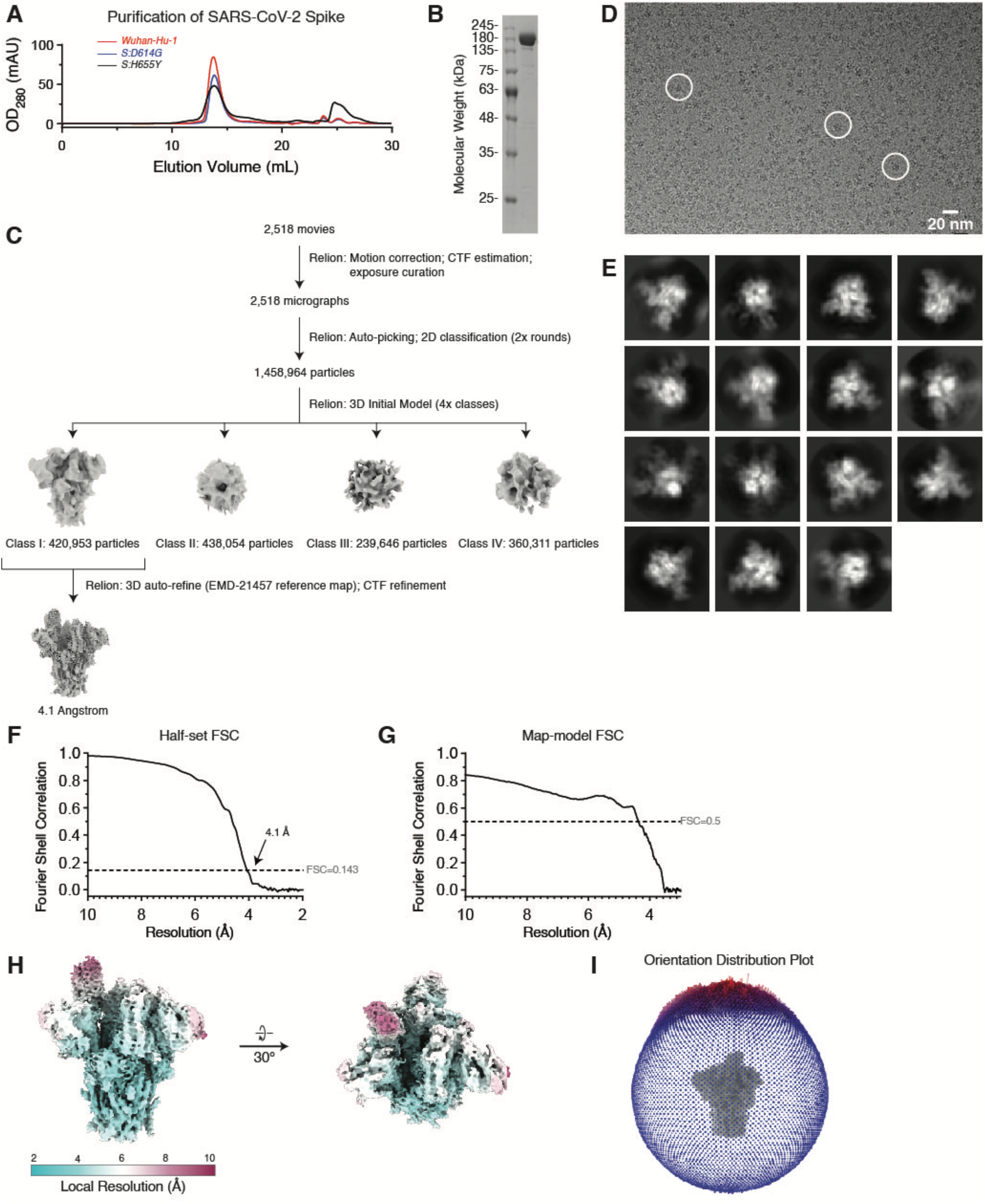
Structural determination of S:H655Y (related to Figure 6). (**A**) Size exclusion column elution profiles of S:H655Y (black line), S:D614G (blue line) and Wuhan-Hu-1 (red line). (**B**) Coomassie-stained SDS-PAGE of S:H655Y shows equivalent, full-length monomeric S proteins. (**C**) Workflow for the data processing of the S:H655Y trimeric complex. (**D**) An exemplary micrograph for the S:H655Y cryo-EM dataset. Individual particles used for downstream data processing are highlighted in white circles. (**E**) 2D clustering of extracted particles. (**F**) Half-set gold-standard Fourier shell correlation (FCS) for the S:H655Y trimeric complex. (**G**) Map-to-model FSC for the S:H655Y trimeric complex. (**H**) Local resolution estimation of the S:H655Y cryo-EM density map. (**I**) Orientation distribution plot for the S:H655Y cryo-EM density map.

**Table 1.** Summary of cryo-EM data collection, three-dimensional reconstruction, and model refinement for S:H655Y.

| **Imaging Parameters and 3D Reconstruction** |  |
| --- | --- |
| Grids | Quantfoil R 1.2/1.3 Au 300 mesh |
| Calibrated magnification | 105,000 |
| Acceleration voltage (kV) | 300 |
| Pixel size (Å) | 0.83 |
| Total dose (e<sup>-</sup>/Å<sup>2</sup>) | 37.22 |
| Exposure time (s) | 1.09 |
| Defocus range (μm) | -0.6 to -1.8 |
| Particles in final 3D reconstruction | 420,953 |
| Resolution ("gold-standard" at FSC 0.143) (Å) | 4.1 |
| **Model refinement** |  |
| Resolution in phenix.real_space_refine (Å) | 4.1 |
| No. atoms: protein | 21,566 |
| RMS deviations: bond lengths (Å) | 0.005 |
| RMS deviations: bond angles (°) | 0.97 |
| MolProbity score | 1.45 |
| EMRinger score | 2.53 |
| Clash score | 4.48 |
| Rotamer outliers (%) | 0.04 |
| Ramachandran angles: favored (%) | 96.51 |
| Ramachandran angles: allowed (%) | 3.49 |
| Ramachandran angles: outliers (%) | 0 |
| **PDB** | 8GB0 |
| **EMDB** | 29910 |

### The receptor binding domain conformation of S:D614G and S:H655Y resembles that of the ancestral Spike in the presence of ACE2

Having demonstrated that S:H655Y promotes an open conformation of the RBD of Wuhan-Hu-1 Spike, we next sought to evaluate the effect of the mutation on Spike conformational dynamics. Specifically, we probed the positioning of the RBD with respect to the N-terminal domain (NTD) using an established single-molecule Förster resonance energy transfer (smFRET) imaging assay^81^. In this assay, soluble, trimeric Spike ectodomain with donor and acceptor fluorophores covalently attached at amino acids 161 and 345, in the NTD and RBD, respectively, were immobilized on a quartz surface and imaged using total internal reflection fluorescence (TIRF) microscopy (Figure 7A). FRET trajectories from individual S:D614 trimers displayed transitions between high-FRET (0.65±0.08) and low-FRET (0.36±0.08) states, indicative of closed and open RBD conformations, respectively, as determined by hidden Markov modeling (HMM) analysis. Trajectories were compiled into histograms to visualize the occupancy of the population of Spike molecules in the two FRET states (Figure 7B). As previously reported, addition of soluble ACE2 or introduction of the S:D614G mutation shifted the equilibrium in favor of the low-FRET open RBD conformation (Figure C-D). In the case of S:D614G, the occupancy of the open RBD conformation was 62 ± 3% (Figure 7D), in agreement with previous structural analysis ^20^. The S:H655Y Spike displayed a similar FRET distribution as S:D614G with 66 ± 2% in the low-FRET state (Figure 7E-F). However, the FRET efficiency of the low-FRET state for S:H655Y shifted downward and became slightly more broad (0.31 ± 0.1). This could indicate a shift in the position of the NTD or inherent variability of the open RBD position. The latter possibility could be reflective of the structural observation that the S:H655Y Spike is only found in a conformation with a single open RBD (Figure 6), whereas multiple open RBDs were seen for Wuhan-Hu-1 and S:D614G^20^. Slight differences in the open RBD position could result from the position of the neighboring RBD.

The HMM analysis also permitted identification of transitions between the FRET states. This analysis is displayed in transition density plots (TDPs) that indicate the relative frequency of transitions between high- and low-FRET states. Addition of ACE2 or the S:D614G protein reduced FRET transitions as compared to the unbound Wuhan-Hu-1 Spike, as previously seen^81^. Most dramatic was the reduction in conformational dynamics seen for S:H655Y (Figure 7J). Further analysis of the kinetics of FRET transitions indicated that the S:H655Y mutation reduced the rate of RBD transition from the closed to the open conformation, and from the open to closed conformation (Figure 7K). This is in contrast to ACE2 binding and S:D614G, which only reduced the rate of transition from the open to the closed conformation. This dramatic reduction in dynamics is consistent with the structural data showing that all S:H655Y Spikes displayed a single conformation (Figure 6).

**Figure 7.**
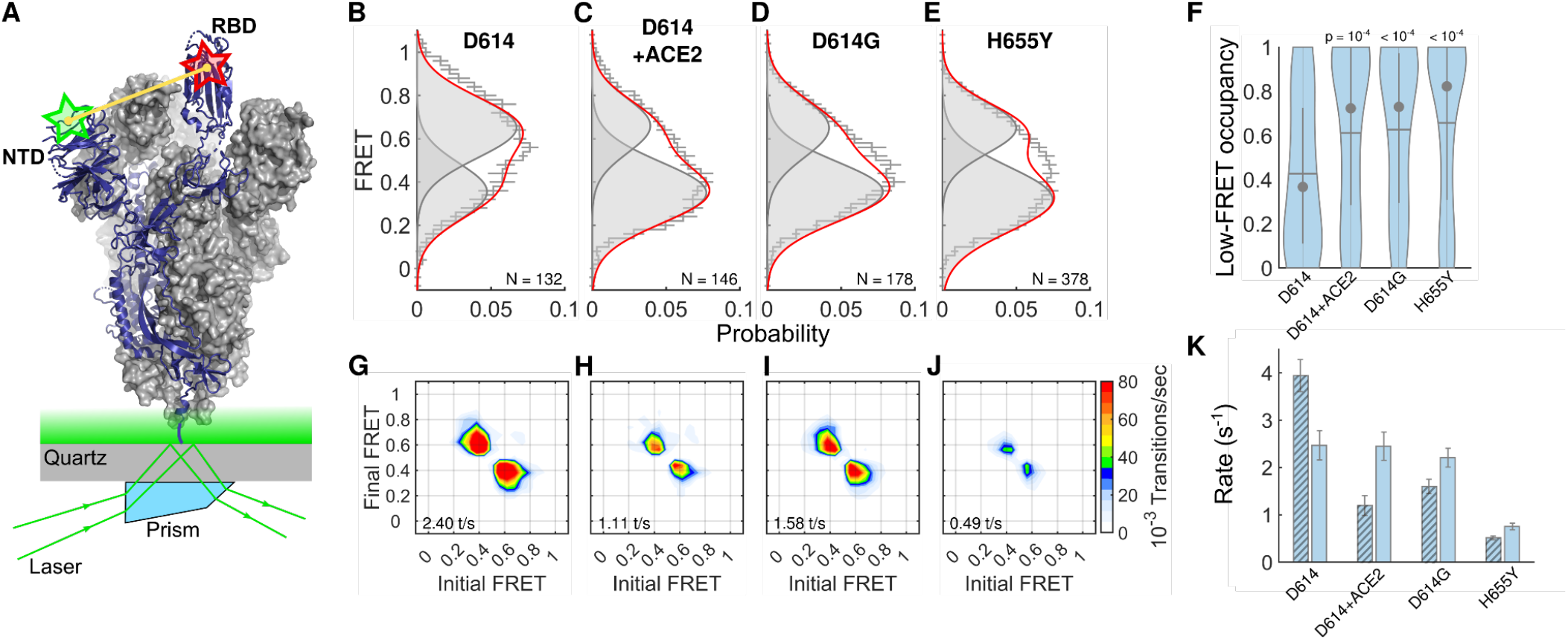
Conformational dynamics of Wuhan-Hu-1, S:D614G, and S:H655Y Spike. **(A)** Experimental setup of smFRET imaging assay that reports on dynamics of RBD with respect to the NTD. Approximate sites of fluorophore attachment are indicated with red and green stars overlaid on a cartoon rendering (blue) of an S protomer, with the unlabeled protomers shown in surface rendering (grey). Soluble S trimers were immobilized on a quartz slide and imaged using TIRF microscopy, as shown. **(B)** FRET histogram compiled from smFRET trajectories acquired from the Wuhan-Hu-1 (S:D614) Spike. Overlaid on the histograms are Guassian distributions, reflecting the mean and width of the high-(closed RBD) and low-FRET (open RBD) states, which were constructed from HMM analysis of the individual smFRET trajectories. N indicates the number of trajectories included in the analysis. **(C-E)** FRET histograms acquired from the indicated Spike complexes, displayed as in (B). **(F)** Violin plots of the occupancies of individual smFRET trajectories in the low-FRET open RBD state. Overlaid on the violin plots are whisker plots indicating the mean (horizontal line), median (circle), and quantiles (vertical lines) of each distribution. Also shown are *p*-values of pairwise comparison with the case of Wuhan-Hu-1 Spike determined by ANOVA. **(G)** TDP indicating the frequency of transitions between the high- and low-FRET states seen in smFRET trajectories acquired on Wuhan-Hu-1 Spike. The total number of transitions per second (t/s) is shown. The frequency of transitions is reflected by the indicated colormap, with red representing the highest frequency. **(H-J)** TDPs for (H) Wuhan-Hu-1 (S:D614) with ACE2 bound, (I) S:D614G, and (J) S:H655Y displayed as in (G). **(K)** The rates of transition between the closed RBD and open RBD conformations for the indicated Spike molecules determined through HMM analysis of individual smFRET trajectories. Rates are presented as the mean +/-standard deviation determined from three independent populations of molecules.

## Discussion

At the conclusion of 2020, SARS-CoV-2 Variants of Concern began to appear with regularity, each time bearing complex combinations of mutations. When first detected, most variants were fully-formed, at the end of long branches on the phylogenetic tree^18^. Intermediate SARS-CoV-2 strains may have gone undetected because complex sets of mutations accumulated during persistent infection within individuals. Sequential sampling from one immunocompromised individual^48^ showed that SARS-CoV-2 infection had initiated with a Spike nearly identical to the parental Wuhan-Hu-1 strain, and that by day 75 post-infection the virus had already accumulated a slew of mutations in common with Alpha, Beta, Gamma, Delta, and Omicron (Figure 1A). Persistent infection would allow SARS-CoV-2 to spread cell-to-cell, without the hurdle of person-to-person transmission, and with minimal exposure to neutralizing antibodies^82^. In this way, SARS-CoV-2 may traverse what would otherwise have been uncrossable fitness valleys^83^, greater numbers of mutations may be sampled, and SARS-CoV-2 variants bearing greater numbers of mutations would have been generated without phylogenetic intermediates being detected in the community^39^.

Notably, at the earliest time point in this persistent SARS-CoV-2 infection, the only S mutation that distinguished it from the ancestral virus was S:D614G^48^, a mutation that was retained at each of the subsequent time points (Figure 1). The experiments here showed that, by day 75 post-infection, when the virus had acquired complex sets of S1 mutations found in Variants of Concern, the cost to infectivity of reverting S:D614G to S:D614 was greater than it would have been in the virus that initiated the infection (Figure 1). If this individual had initially been infected with the ancestral SARS-CoV-2 that bears S:D614, the S mutations found at day 75 may have been incapable of sustained replication and the infection would have been cleared sooner.

The effect of S:D614G on the S sequences from the persistent infection (Figure 1), along with the fact that Variants of Concern were only detected after S:D614G had gone to near fixation, spurred assessment of the importance of S:D614G for the infectivity of Variants of Concern. Each variant was more S:D614G-dependent than the ancestral virus (Figures 2, 3, and 4). For example, reversion of S:D614G on the Alpha background decreased infectivity more than 70-fold, an effect that was more than ten-fold greater than for the ancestral Wuhan-Hu-1 (Figure 2). Thus, while S:D614G increased the infectivity of Wuhan-Hu-1 Spike, this effect was greatly amplified via epistatic interactions in the context of the mutations present in Variants such as Alpha^84^.

On the background of the Wuhan-Hu-1 Spike, S:D614G modestly increased the amount of S1 that was retained on virions (Figure 2). When the effect of S:D614G was tested on the background of Spikes from different variants, the magnitude of Spike stabilization increased in parallel with the increase in infectivity (Figure 2, 3, and 4). This indicates that, by stabilizing the mature Spike protein on virions, S:D614G acts epistatically to facilitate acquisition of other S mutations, including K417N, E484K, and N501Y (Supplementary Figure 1). In contrast to S:D614G, which did not increase ACE2-binding affinity and did not confer escape from neutralizing antibodies^19, 20^, the latter mutations increased affinity for ACE2 and conferred the ability to escape from neutralizing antibodies^85–88^. Similar evolutionary pathways, in which mutations of relevance to immune escape were detrimental to replication unless preceded by stabilizing mutations, have been reported for influenza A virus^89^.

Variants that circulated in the absence of S:D614G were sought, with the idea that they might harbor mutations with similar phenotypes to S:D614G, and thereby facilitate attempts to understand how convergent traits evolve in SARS-CoV-2^90^. Three exceptional variants were identified and each was shown to be dependent upon either S:Q613H or S:H655Y (Figure 3). All three mutations, S:Q614H, S:D614G, and S:H655Y, were then found to stabilize mature Spike protein on virions (Figures 2, 3, and 4), consistent with their selection via a common molecular mechanism. These apparent examples of convergent evolution strengthen structure-function correlations concerning the molecular mechanism of action of S:D614G.

Gamma and Omicron variants have both S:D614G and S:H655Y (Supplementary Figures 1 and 2), and the infectivity of Gamma and Omicron (BA.1) was dependent upon both S:D614G and S:H655Y (Figure 3 and 4). That being said, the relative influence of S:D614G and S:H655Y was highly-dependent upon the Spike amino acid context. Infectivity of Wuhan Hu-1 Spike was increased to a similar extent by S:D614G and S:H655Y, while the Gamma variant was more dependent upon S:D614G than on S:H655Y (Figures 2 and 3). In contrast, the Omicron (BA.1) variant was more dependent upon S:H655Y than on S:D614G (Figure 3 and 4) and when S:H655Y was reverted to the ancestral sequence on the A.30 Spike, S:D614G was incapable of rescuing infectivity.

SARS-CoV-2 is the only *Sarbecovirus* known to have a polybasic S1/S2 cleavage site, a feature that contributes to the tissue tropism, pathogenicity, and immune escape of this deadly human pathogen^74^. S:D614G and S:H655Y have rarely been detected in other *Sarbecoviruses* and, after S:D614G had become fixed in circulating SARS-CoV-2, there was a steady stream of variants bearing mutations with increased S1/S2 processing efficiency^91^. These observations suggested that the replication advantage of S:D614G and S:H655Y might only be evident in the presence of the polybasic S1/S2 cleavage site, and that these mutations would promote infectivity by maintaining non-covalent association of S1 with the virion, in the face of more efficient S protein cleavage.

To test this hypothesis, dependency on S:D614G or S:H655 was evaluated here under conditions in which S1/S2 processing efficiency was either decreased or increased using orthogonal genetic and pharmacologic perturbations. S:D614G had no effect on the infectivity of SARS-CoV-1, a virus that does not have a polybasic S1/S2 cleavage site (Figure 5). When the SARS-CoV-2 S1/S2 cleavage site was introduced into the SARS-CoV-1 Spike it became as S:D614G-dependent as was the SARS-CoV-2 Wuhan Hu-1 Spike (Figure 5). Conversely, disruption of the polybasic S1/S2 cleavage site in the SARS-CoV-2 Wuhan-Hu-1 Spike, or production of Spike in the presence of furin protease inhibitors, rendered infectivity independent of S:D614G (Figure 5). This effect was more dramatic with the highly S:D614G-dependent SARS-CoV-2 Alpha Spike; disruption of the cleavage site increased infectivity of the S:D614 revertant 70-fold (Figure 5). Similarly, the magnitude of Wuhan-Hu-1 Spike dependence on S:D614G or S:H655Y was greatly increased in the presence of S:P681R, a mutation found in the Delta variant that increases cleavage efficiency^68^. Finally, when reverse genetic Omicron (BA.1) was propagated in tissue culture, the S1/S2 cleavage site was retained in the wild type and in the single revertant mutants (either to S:D614 or to S:H655), but it was already deleted from the double reversion mutant (S:D614/H655) within one passage in tissue culture (Figure 6L). In all cases mentioned above, disruption of the polybasic S1/S2 cleavage site maintained virion-associated S1 protein levels on the virion as did S:D614G and/or S:H655Y (Figure 5).

Single-particle CryoEM revealed that both S:D614G^20^ and S:H655Y (Figure 6) dramatically shift the probability that the SARS-CoV-2 Spike receptor binding domains populate the open conformation required for ACE2-binding (Figure 6B). Separately, smFRET showed that Spikes bearing either S:D614G or S:H655Y spontaneously mimic a conformation that is induced in the presence of ACE2 (Figure 7). Given that ACE2-binding is precluded when Spike RBD is in the closed conformation^78, 79^, the smFRET data is consistent with both mutations promoting a more open RBD conformation. The convergence of the findings from these orthogonal biophysical methods buttress the conclusion that S:D614G and S:H655Y promote an open RBD confirmation.

To pinpoint the effect of each mutation in these structural and biophysical studies, S:D614G and S:H655Y were assessed as individual mutations on an isogenic Wuhan Hu-1 background. Given the much greater importance of S:D614G for infectivity on the background of the Alpha Variant Spike (Figure 2), one might expect an even greater shift to open conformations of the receptor binding domain, but structural studies of the S:D614G revertant on the Alpha Spike background were precluded by instability of the soluble trimer. Nonetheless, a shift to higher frequency open conformations has been seen with Spikes from the Gamma and Omicron BA.1 variants^92–94^. The greater openness of these variant Spikes suggests that S:D614G and S:D655Y promote more open conformation in the context of these complex sets of mutations.

Though the ACE2-binding site is only accessible on the Spike RBD open conformation, which is more common with S:D614G and S:H655Y, the increased infectivity of these mutants is not explained by increased affinity for ACE2 (Supplementary Figure 3). Rather, increased infectivity is more likely a result of greater Spike stability on virions following cleavage at the S1/S2 junction. The stability conferred by S:D614G and S:H655Y is evident in western blots of both pseudovirions (Figures 2, 3, and 5), and *bona fide* virions (Figure 4), after enrichment by acceleration in an ultracentrifuge. This increased stability is also reflected in the smFRET data which showed reduced conformational dynamics of Spike trimers bearing either S:D614G and S:H655Y (Figure 7).

The net effect of S:D614G and S:H655Y on Spike stability and more open conformation is the same, though the atomic details may differ. The structural basis for the increased stability of S:D614G was previously attributed to the ordering of the 630 loop, which expanded hydrophobic contacts with the CTD2 domain^75^. Close examination of the atomic model of Spike bearing S:H655Y shows hydrophobic stacking of S:H655Y with the sidechain of S:F643 (Figure 6C). Perhaps more important for noncovalent stabilization of S1 on the virion is electrostatic contact between S:H655Y in S1 and S:T696 in S2 (Figure 6D). Following cleavage, this contact could strengthen the S1/S2 interaction, leading to reduced S1 shedding, and a more stabilized S protein complex on the virion. In addition, as the RBD shifts from closed to open conformation, the distances between these two pairs of residues do not change in S:D614G and S:H655Y, whereas both increase by ∼0.5 Å in the Wuhan Hu-1 Spike. The bigger shifts with Wuhan Hu-1 Spike suggest how S1 is lost more readily when the RBD shifts from closed to open conformation, and why Spike trimers with more open conformation are more readily detected with S:D614G and S:H655Y.

## Acknowledgements

We thank members of the Munro, Shi, Shen, and Luban labs for technical assistance and helpful discussions. This work was supported by grants from the Massachusetts Consortium on Pathogen Readiness to J.L. and K.S., Project PRJ-6691 from the Coalition for Epidemic Preparedness Innovations (CEPI) to J.L., NIH grants R37AI147868 and R01AI148784 to J.L., a Sara Elizabeth O’Brien Fellowship Award/King Trust to L.Y., and NIH grant R01GM143773 to J.B.M. P.-Y.S. was supported by NIH contract HHSN272201600013C and awards from the Sealy & Smith Foundation. We thank the global research community for rapid sharing of SARS-CoV-2 sequence data and the organizers of the platforms that make this data accessible and interpretable, including: https://nextstrain.org/, https://www.gisaid.org/, https://www.pango.network/, https://outbreak.info/, https://covdb.stanford.edu/, and https://covariants.org/.

## Materials and Methods

### Plasmids and DNA engineering

The plasmids used here were either previously described or generated using standard cloning methods. The full list of plasmids used here is provided in Table 2. Previously engineered plasmids are listed with their source. All newly engineered plasmids are listed with Addgene accession numbers (https://www.addgene.org/Jeremy_Luban/), where they are publicly available with their full sequences.

### Cell culture

All cells used here were obtained from the American Type Culture Collection (https://www.atcc.org). Human cells lines were cultured in humidified incubators with 5% CO_2_ at 37° C, and monitored for mycoplasma contamination using the Mycoplasma Detection kit (Lonza LT07-318). HEK-293T cells (CRL-11268) were cultured in DMEM supplemented with 10% heat-inactivated FBS, 1 mM sodium pyruvate, 20 mM GlutaMAX, 1× MEM non-essential amino acids, and 25 mM HEPES, pH 7.2.

### Production of single-cycle lentiviruses pseudotyped with SARS-CoV-2 Spike

24 hrs prior to transfection, 6 × 10^5^ HEK-293T cells were plated per well in 6 well plates. All transfections used 2.49 µg plasmid DNA with 6.25 µL TransIT LT1 transfection reagent (Mirus, Madison, WI) in 250 µL Opti-MEM (Gibco). Single-cycle HIV-1 vectors pseudotyped with the indicated SARS-CoV-2 Spike constructs, were produced by transfection of HIV-1 pNL4-3ΔenvΔvpr luciferase reporter plasmid (pNL4-3.Luc.R-E-; NIH AIDS Reagent Program, Division of AIDS, NIAID, NIH: from Dr. Nathaniel Landau; ARP Cat #3418) in combination with the indicated Spike expression plasmid, at a ratio of 4:1. All lentivirus preparations were normalized with a quantitative, in-house, exogenous reverse transcriptase assay as previously described^20^.

### Single-cycle lentivirus infectivity assay

16 hours prior to transduction, HEK-293T cells stably expressing ACE2/TMPRRS2 as previously described^20^ were plated at 3 x 10^4^ per well. Cells were incubated in virus-containing media for 16 hours at 37°C after which fresh media was added to cells. 48 hours after transduction cells were assessed for luciferase activity using the Promega Steady-Glo system (Promega Madison, WI).

### Western blot analysis

Tissue culture media and cell lysate were collected 60 hours after transfections to produce lentivectors. Supernatant containing Spike-pseudotyped particles was layered on a 20% sucrose cushion in PBS and spun at 110,000 x g at 4°C for 2 hrs. The pellet was washed once with ice cold PBS and resuspended in 15 uL of 2x SDS gel loading buffer. After removal of supernatant, transfected cells were lysed in 300 uL 2x SDS-PAGE loading buffer. Protein preps were boiled for 5 minutes and then separated by SDS-PAGE on a 10-20% Tris-Gycine gel (BioRad). Proteins were electro-transferred from gels to nitrocellulose membranes, which were blocked for an hour with Licor Blocking Buffer and detected with the indicated antibodies.

### Reverse genetic system to construct Omicon (BA.1) SARS-CoV-2 viruses bearing S:D614G/H655Y (wild-type), S:D614/H655Y, S:D614G/H655, or S:D614/H655

An infectious cDNA clone of Omicron (BA.1) SARS-CoV-2 was generated using PCR-based mutagenesis, as previously described^66, 67^. To construct S mutations encoding reversions to the Wuhan-Hu-1 residues, either S:D614, S:H655, or S:D614D/H655, nucleotide substitutions were introduced by oligonucleotide mutagenesis into the pcc1-CoV-2-BA1-F567 subclone containing the SARS-CoV-2 BA.1 S gene. All primers used for site-directed mutagenesis are listed in Table 2. The resulting plasmids were validated by Sanger sequencing. The full-length infectious cDNA clone of SARS-CoV-2 was assembled by in vitro ligation of three contiguous cDNA fragments following the previously described protocol ^67, 68^. In vitro transcription was performed to synthesize full length viral genomic RNA. The RNA transcripts were electroporated in Vero E6 cells expressing TMPRSS2 (purchased from SEKISUI XenoTech, LLC) to recover the mutant viruses. Viruses were rescued post 2-4 days after electroporation and served as P0 stock. P0 stock was passed once more on Vero E6 cells expressing TMPRSS2 to produce P1 stock.

The S1 gene was sequenced from all P1 stock viruses to confirm the presence of the engineered mutations. P1 stock was titrated by plaque assay on Vero E6 cells expressing TMPRSS2 and used for subsequent experiments. All virus preparation and infections were carried out at biosafety level 3 (BSL-3) facility at the University of Texas Medical Branch at Galveston.

### RNA extraction, RT-PCR, and cDNA sequencing

Cell culture supernatants or P1 virus cultures were mixed with a five-fold excess of TRIzol™ LS Reagent (Thermo Fisher Scientific, Waltham, MA). Viral RNAs were extracted by Direct-Zol™ RNA miniprep Plus Kit (ZYMO RESEARCH) according to the manufacturer’s instructions. The extracted RNAs were dissolved in 50 μl nuclease-free water. For sequence validation of mutant viruses, 5 µl of RNA samples were used for reverse transcription by using the SuperScript IV First-Strand Synthesis System (Thermo Fisher Scientific) with primers flanking the spike gene. The resulting DNAs were gel extracted by QIAquick gel extraction Kit, and the genome sequences were determined by Sanger sequencing at GENEWIZ (South Plainfield, NJ).

### Plaque assay

Approximately 1×10^6^ Vero E6 cells expressing TMPRSS2 were seeded to each well of 6-well plates and cultured at 37°C, 5% CO_2_ overnight. Virus was serially diluted in DMEM with 2% FBS and 200 µl diluted viruses were transferred onto the cell monolayers. The viruses were incubated with the cells at 37°C with 5% CO_2_ for 1 h and then replaced with 2 ml of overlay medium containing DMEM with 2% FBS, 1% penicillin/streptomycin, and 1% sea-plaque agarose (Lonza, Walkersville, MD). After 2 days of incubation, plates were stained with neutral red (Sigma-Aldrich, St. Louis, MO, USA) and plaques were counted on a light box.

### Primary human airway cultures infection

The EpiAirway system is a primary human airway 3D mucociliary tissue model consisting of normal, human-derived tracheal/bronchial epithelial cells (https://www.mattek.com/products/epiairway/). Wild-type SARS CoV-2 Omicron (BA.1) was mixed in a 3:1 ratio with each BA.1 revertant virus (S:D614/H655Y, S:D614G/H655, or S:D614/H655) based on the PFU/ml titer of the virus. Mixed viruses were inoculated onto the culture at a total MOI of 0.2 in DPBS. After 2 h infection at 37°C with 5% CO2, the inoculum was removed, and the culture was washed three times with DPBS. The infected epithelial cells were maintained with medium in the basal well and no medium in the apical well. The infected cells were incubated at 37°C, 5% CO2. From day 1 to day 3 post-infection, 300 μl of DPBS was added onto the apical side of the airway culture and incubated at 37°C for 30 min to elute progeny viruses. 200 μl of this sample was mixed with 800 μl TRIZOL-LS. On day 3, after collecting progeny viruses, 300 μl of TRIZOL was added to the cells and after 5 min incubation, 200 μl TRIZOL was collected for isolating cellular RNA.

### Next generation sequencing (NGS) to determine relative replicative fitness

Viral RNA samples from the three competition experiments, wild-type BA.1 (S:D614G/H655Y) versus either S:D614/H655Y, S:D614G/H655, or S:D614/H655Y, were used as template in a one-step RT-PCR reaction with the SuperScrip IV One-Step RT-PCR kit (Invitrogen, Carlsbad, CA, USA) and primers that amplify coding sequences for Spike amino acids 614 and 655 (Table 2). cDNA synthesis was for 10 min at 50°C with pre-denaturation for 2 min at 94°C and 25 cycles of denaturation at 94°C for 10 s, annealing at 58°C for 10 s, and extension at 72°C for 15 s. A final extension was at 72°C for 5 min. The PCR products were enriched with a QIAquick PCR Purification kit (Qiagen, Germantown, MD) according to the manufacturer’s protocol. Dual-indexed adapter sequences (New England BioLabs, Ipswich, MA) were added with 5 cycles of PCR. Samples were pooled and sequenced on an Illumina MiniSeq Mid-Output flow cell with the paired-end 150 base protocol. Reads were filtered for the highest quality Q-scores of 37^95^ at the coding sequences for Spike residue 614 (GGT or GAT) and 655 (TAT or CAT). The input and output ratios of each pair of competing viruses were calculated and used to determine the relative replicative fitness^68^.

### Virion purification and western blotting

Vero E6 cells (Figure 5K), or Vero E6 cells bearing TMPRSS2 (Figure 4C), were infected with SARS-CoV-2 Omicron BA.1 S:D614G/H655Y (wild-type), S:D614/H655Y, S:D614G/H655, or S:D614/H655S viruses at an MOI of 0.1. At 48 h post-infection, the culture medium was collected and centrifuged to clear the cell debris. Cell supernatant was mixed with polyethylene glycol (PEG) at a final concentration of 10% and mixed continuously for 30 min at RT and then accelerated in an Eppendorf Centrifuge 5810R at 3,197 x g for 10 min to pellet the virions. The virus pellet was washed once with 70% ethanol. The pellet was suspended in 2x Laemmli sample buffer containing β-mercaptoethanol (BME) and analyzed by Western blot on a 4–20% gradient SDS-PAGE gel. The following antibodies were used to detect the S1 (Sino biological #40591-T62), S2 (Invitrogen #MA5-35946), and N (Novus #NB100-56576) proteins of SARS CoV-2.

### Purification of soluble SARS-CoV-2 S

FreeStyle 293-F cells were cultured in SMM-293 TII serum-free media (SinoBiological) and maintained at 37°C with shaking and 8% CO2 and 80% humidity. 500 μg of plasmid encoding His-tagged S:H655Y was transfected into 400 mL of 293 FreeStyle cells at 5×10^5^ cells/mL. 72 h later, the media was filtered using a 0.45 μm filter, and furthermore applied to Ni-NTA resin (QIAGEN). The resin was collected and washed with PBS supplemented with 20 mM imidazole to mitigate non-specific binding of proteins. Lastly, soluble S:H655Y was eluted using PBS supplemented with 200 mM imidazole, and then further purified using a Superose 6 gel-filtration column (GE healthcare), in buffer containing 25 mM Na-HEPES (pH 7.4) and 150 mM NaCl.

### Measurement of ACE2 binding kinetics

Kinetics of SARS-CoV-2 Spike binding to ACE2-Fc was assessed by surface plasmon resonance using a Protein A sensor kit and a Nicoya OpenSPR instrument (Nicoya, Kitchener, ON, Canada). Protein A was diluted in 10 mM sodium acetate pH 5.0, immobilized on a high-capacity carboxyl sensor chip, and the chip was blocked with 1M ethanolamine. 5 µg/mL of hACE2-Fc (AcroBiosystems, Newark, DE, USA) was injected at a 20 µL/min flow rate for 300 sec. with a minimum response of 100 resonance units (RU). Soluble, stabilized, trimeric Wuhan Hu-1, S:D614G, and S:H655Y Spikes proteins, produced in FreeStyle 293-F cells as described above, were dialyzed in a PBS-T-BSA running buffer (phosphate buffered saline [PBS] pH 7.4, 0.05% v/v Tween 20, and 1% w/v bovine serum albumin [BSA]). Binding kinetics was evaluated by injecting 2-fold serial dilutions (6.25-100 nM) of each soluble SARS-CoV-2 Spike trimer protein. Spike analytes were injected at a flow rate of 50 µL/min for 120 sec (association phase), followed by a 270 sec dissociation phase. After each analyte injection, full regeneration was achieved by injecting 10 mM glycine HCl pH 3.0 at a flow rate 150 µL/min for 40 sec. Following regeneration, and before the next analyte injection, 5 µg/mL of hACE2-Fc was injected at the rate and duration described before. After each step in the cycle, PBS-T-BSA was injected for 300 sec to clean the surface prior to the subsequent injection. Sensorgrams were double reference subtracted before the kinetics analyses were performed, as described^96^. The kinetic analysis was performed using TraceDrawer software v.1.9.1 (Nicoya, Kitchener, ON, Canada) and kinetic parameters were obtained by fitting the data to a 1:1 binding model. The association rate constant (*k_a_*) was determined by fitting the analyte binding at each concentration, and the dissociation rate constant (*k_d_*) was determined by fitting the change in the binding response during both the association and the dissociation phases. The *K_D_* (equilibrium dissociation constant) was calculated from the ratio between the *k_a_* and *k_d_* constants.

### Cryo-EM sample preparation and data collection

3.5 μL of enriched S:H655Y, at 2.0 μg/μL, was deposited on an UltrAuFoil R1.2/1.3 300 mesh grid that had been previously glow discharged for 30 s in a GloQube Plus Glow Discharge System. Following deposition, plunge freezing was performed with a Vitrobot Mark IV using a blot force 3 and 3 s blot time at 95% humidity and 10°C. Following, frozen grids were then imaged using a Thermo Scientific KriosRx

Cryo-Transmission Electron Microscope operated at 300 kV. Our images were recorded using a Falcon 4 Direct Electron Detector. Our movies were monitored using EPU 2 software. In total, 2,518 micrographs were collected using a nominal defocus range of 0.6 to 1.8 μm, at a nominal magnification of 105,000x and a pixel size of 0.83Å. In order to mitigate the preferred orientation issue common throughout cryo-EM studies of the S protein, a tilt-angle of 30 degrees was used for all data collected. We set the dose rate to 37.2 electron/Å^2^.

### Cryo-EM data processing

Processing of the collected micrographs was carried out in Relion 4.0.0. First, movie frame alignment was completed using the motionCorr2 implementation. Following motion-correction, patch-CTF estimation, particle picking, and numerous rounds of 2D classification resulted in 2D projections with recognizable features that were reminiscent of previous structural characterizations of the S protein. Upon completion of 2D classification, 3D initial model building (4 classes) was carried out using the published structure, EMD-21457, as the initial model in Relion 4.0.0. From this, only one of the classes resembled the Spike protein, whereas the three additional classes resembled junk particles. Ensemble 3D auto-refinement followed by CTF correction of 420,953 particles resulted in a 4.1Å map according to “gold standard” Fourier shell correlation of 0.143. The final map was deposited with the accession code EMD-29910.

### Building and validation of structural models

Atomic models were prepared using Coot. Previously deposited structural models of the Wuhan-Hu-1 Spike were used as an initial starting point for model building (PDB: 6VYB). Following, PHENIX was utilized to perform real-space refinements. To assess the validity of our structural model, MolProbity was used to evaluate the model geometries. Corrected Fourier shell correlation curves were calculated using the refined atomic model and the cryo-EM density map. The coordinates were deposited with the accession code 8GB0. To validate the structural model, we assessed the “Gold Standard” FSC curve, as well as the map-to-model FSC curve.

### smFRET imaging

The SARS-CoV-2 SDTM from the Wuhan-Hu-1 strain (residues Q14–K1211) with SGAG substitution at the furin cleavage site (R682 to R685), and proline substitutions at K986 and V987 was expressed and purified as previously described^81^. The D614G mutation was introduced as described^81^. The H655Y mutation was introduced through overlap-extension PCR using the primers S_XhoI_1, S_H655Y_2, S_H655Y_3, and S_ClaI_4 (Table 2). Mutants were confirmed through Sanger sequencing (GENEWIZ^®^, Cambridge, MA, USA). To enable fluorophore attachment for smFRET imaging, SDTM trimers were formed with a single protomer containing A4 peptide insertions in the RBD and NTD domains at positions 161 and 345, respectively, as described^81^. Briefly, SDTM hetero-trimers for smFRET experiments were expressed by co-transfection of ExpiCHO-S cells (Thermo Scientific, Waltham, MA, USA) with both the untagged SDTM construct and the corresponding 161/345A4-tagged SDTM plasmid at a 2:1 ratio. After their purification by affinity chromatography using Ni-NTA agarose beads (Invitrogen, Waltham, MA, USA) and size exclusion chromatography (SEC; Cytiva, Marlborough, MA), SDTM hetero-trimers were labeled by overnight incubation at room temperature with coenzyme A (CoA)-conjugated LD550 and LD650 fluorophores (Lumidyne Technologies, New York, NY, USA) and Acyl carrier protein synthase (AcpS). Labeled trimers were then purified away from unbound fluorophore and AcpS by a second round of SEC. Aliquots were stored at -80°C until use.

The 6X-His tagged labeled SDTM spikes were immobilized on streptavidin-coated quartz microscope slides by way of Ni-NTA-biotin and imaged using wide-field prism-based TIRF microscopy as described^81, 97^. smFRET data were collected using Micromanager v2.0 at 25 frames/s (micro-manager.org,^98^). All smFRET data were processed and analyzed using the SPARTAN software package in Matlab (Mathworks, Natick, MA)^99^. smFRET traces were identified according to following criteria: duration of smFRET trajectory exceeded five frames, correlation coefficient calculated from the donor and acceptor fluorescence traces ranged between –1.1 and -0.1, and signal-to-noise ratio was greater than 8. Traces that fulfilled these criteria were then verified manually. Traces from each of three technical replicates were compiled into FRET histograms and the mean probability per histogram bin ± standard error were calculated. Traces were idealized to a three-state HMM (two non-zero-FRET states and a 0-FRET state) and the transition rates were optimized using the maximum point likelihood (MPL) algorithm^100^ implemented in SPARTAN. The rates reported in Figure 7K reflect the mean ± standard error determined from three technical replicates. The three-state model was selected by comparing the Akaike information criterion (AIC) across multiple different models with a range of state numbers and topologies as previously described ^81^. The idealizations from the total population of traces analyzed were used to construct Gaussian distributions, which were overlaid on the FRET histograms to visualize the results of the HMM analysis. These idealizations were also used to calculate the occupancies of each FRET trajectory in the 0.65- and 0.35-FRET states. The distributions in occupancies were used to construct violin plots in Matlab, as well as calculate median occupancy, mean occupancy, 25th and 75th quantiles, and standard errors, as displayed in Figure 7F. Statistical significance measures (p-values) were determined by one-way ANOVA in Matlab. This analysis displays the full breadth of dynamic behavior across the total population of traces analyzed. The total number of traces analyzed was sufficient to ensure minimally 85% statistical power during comparisons ^81^.

### Quantitation and Statistical Analysis

GraphPad Prism 8.4.3 was used to analyze the infectivity data using a ratio paired t test. In these experiments, all values shown are the mean with standard deviation, with the actual calculated two-tailed *P* value indicated on the figure. In Figure 4, simple linear regression was performed to analyze the statistical significance of the RNA ratio at a given time-point versus the input RNA ratio for the competition experiment.

---

**Supplementary Table 2.**
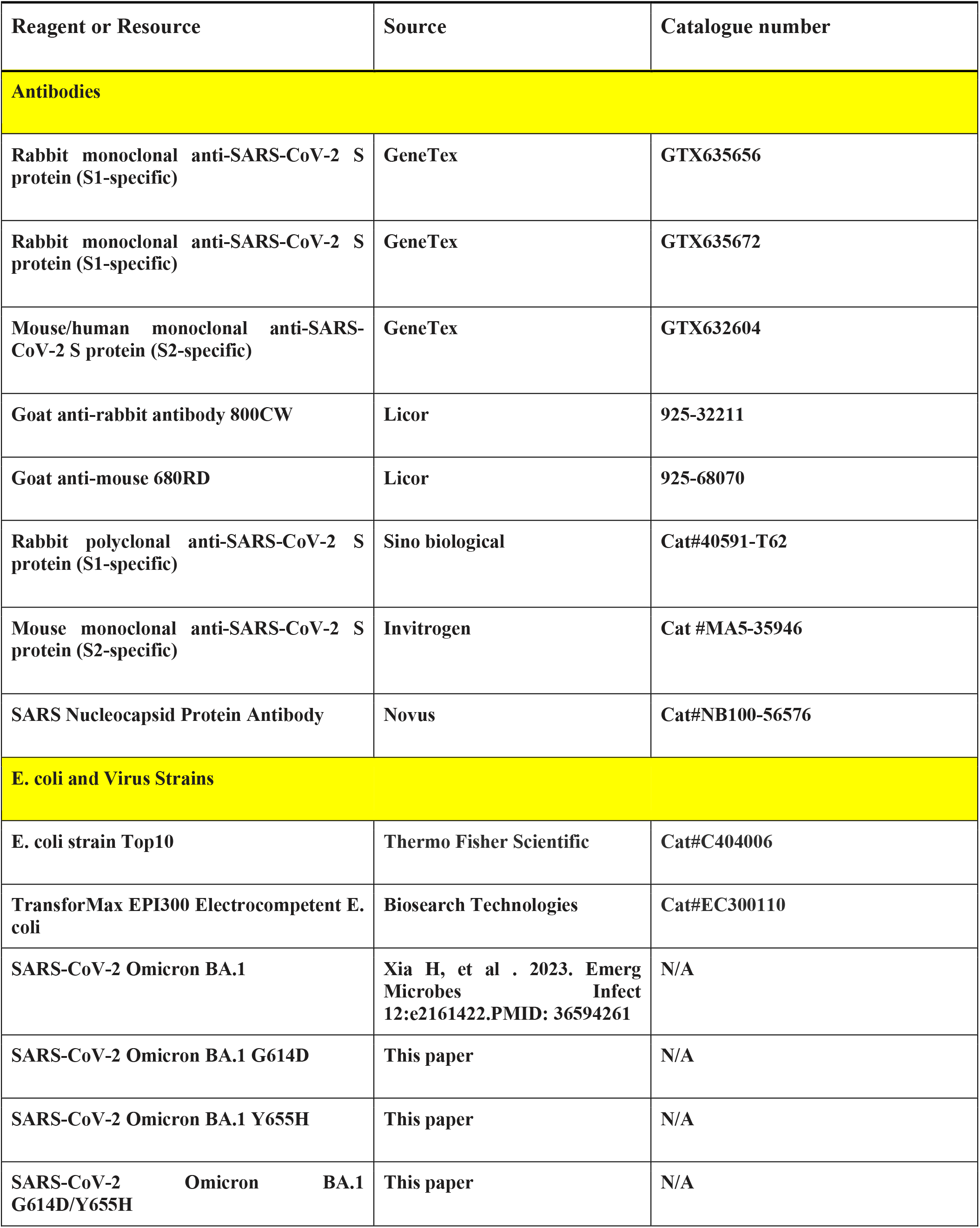
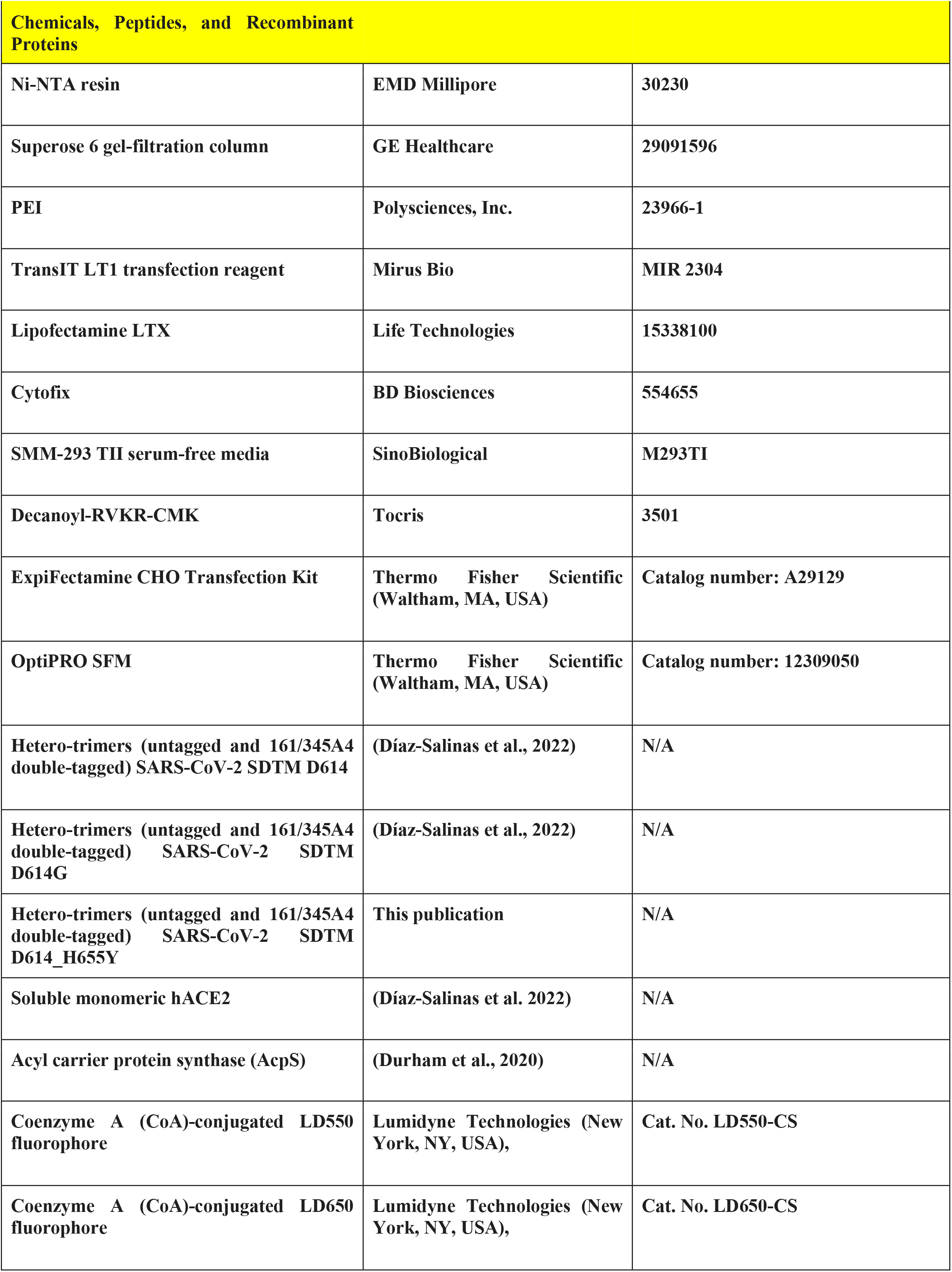
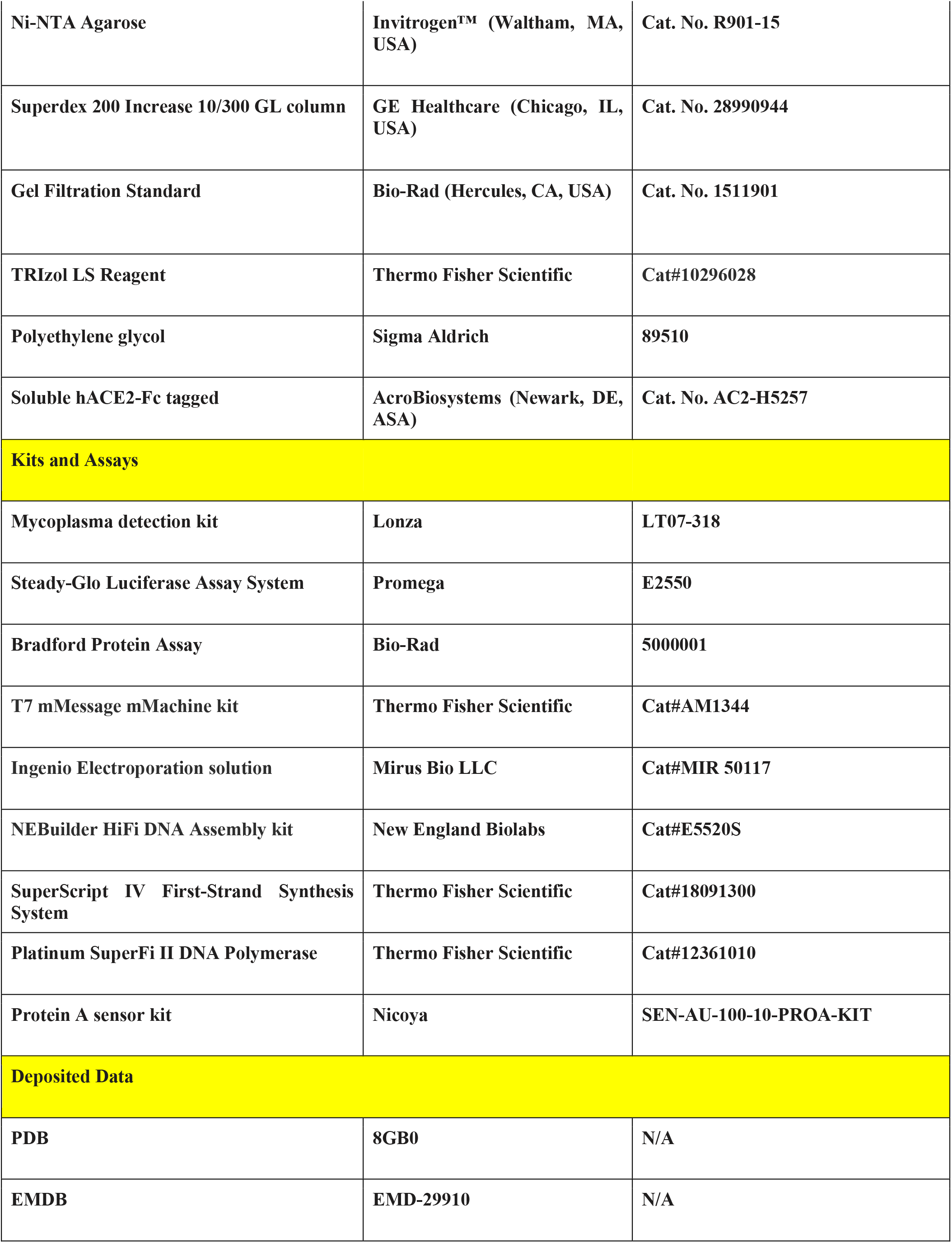
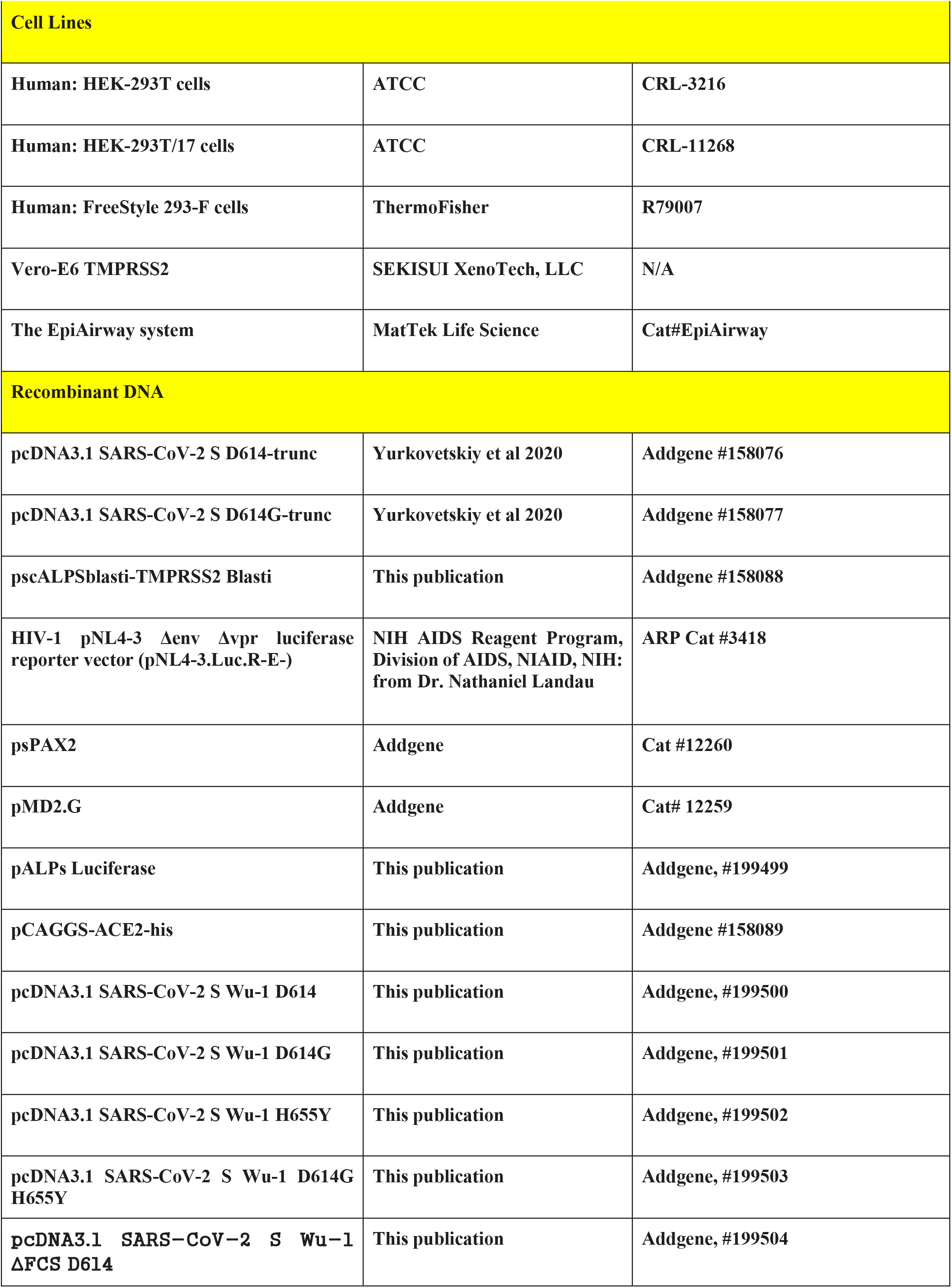
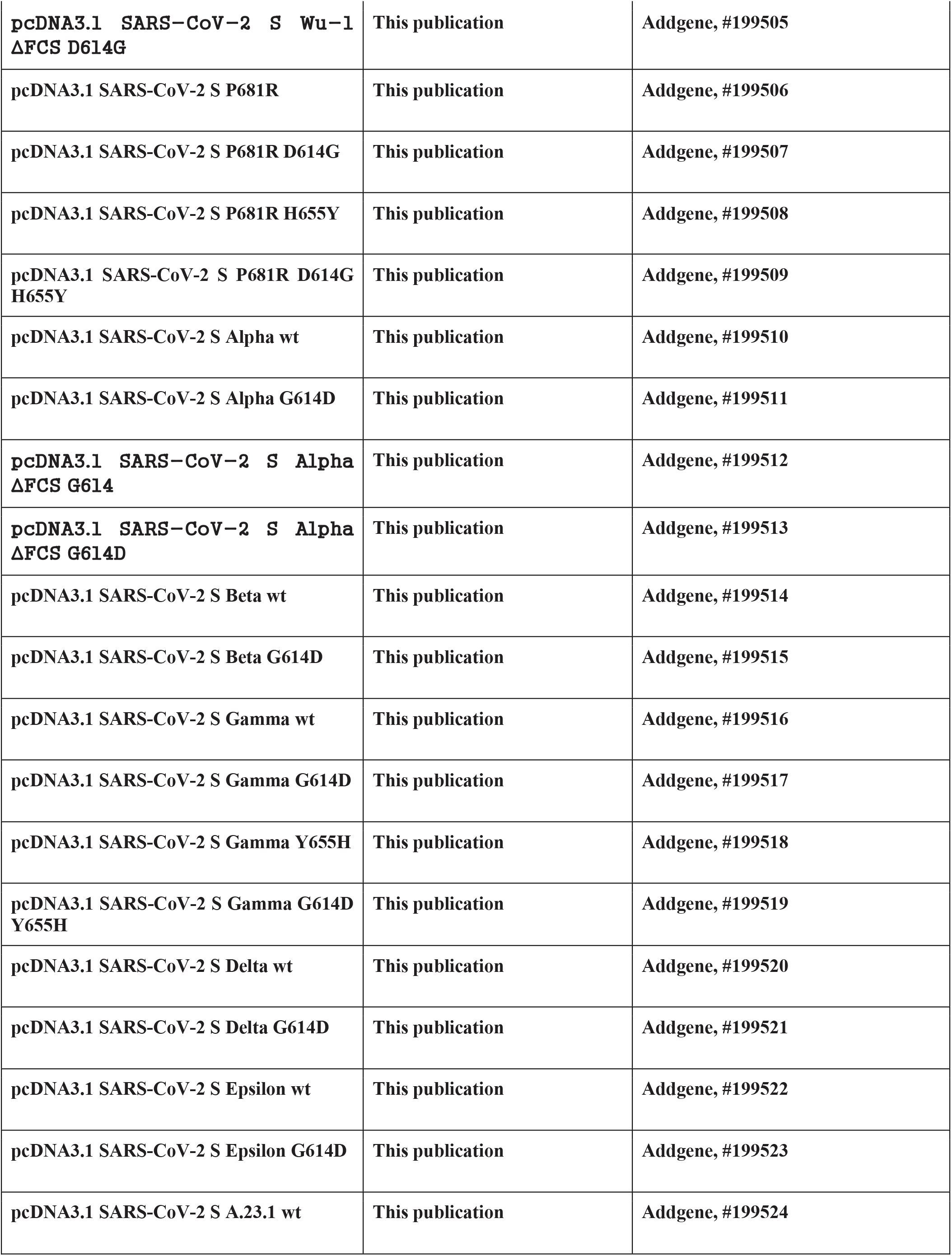
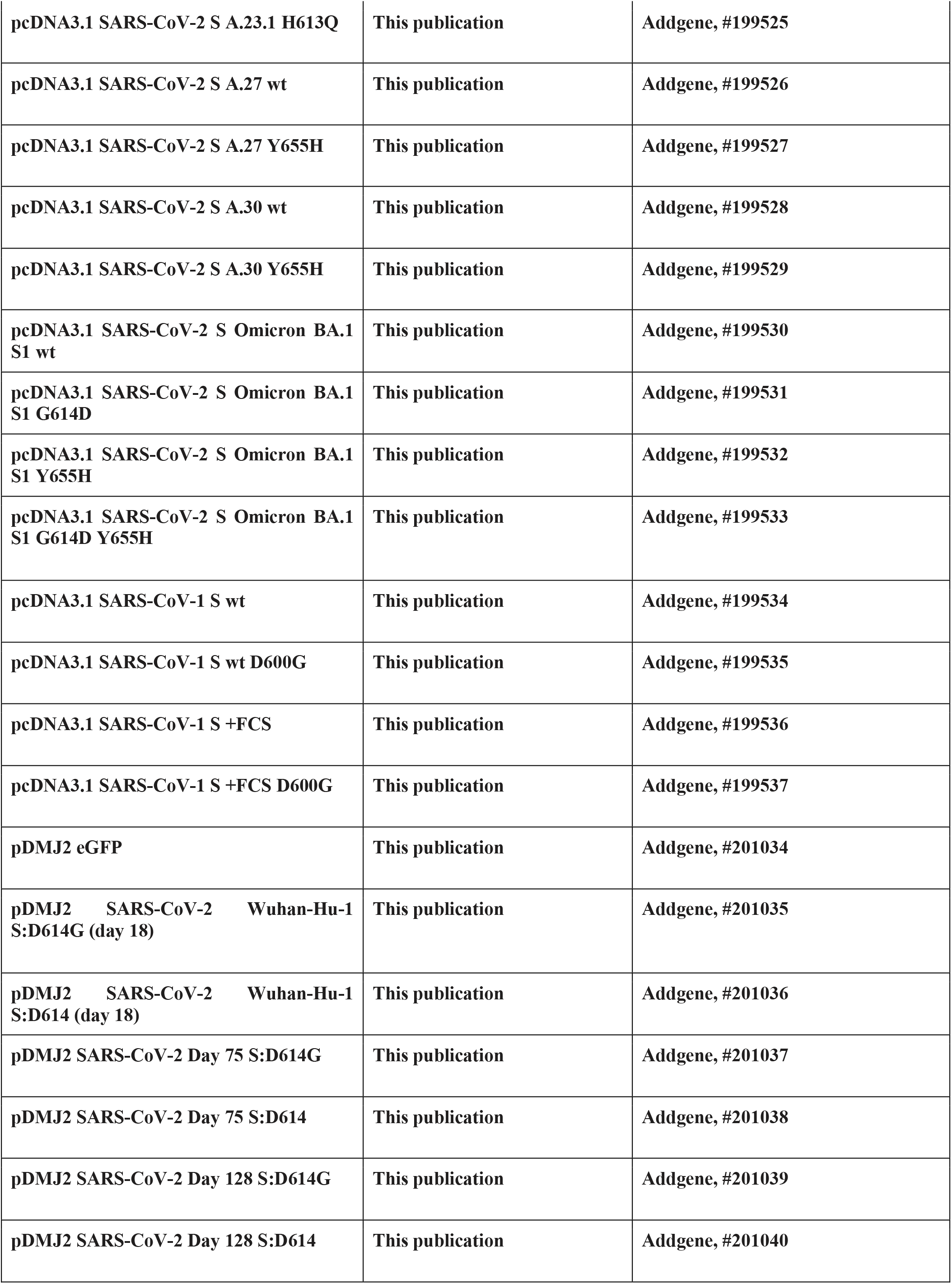
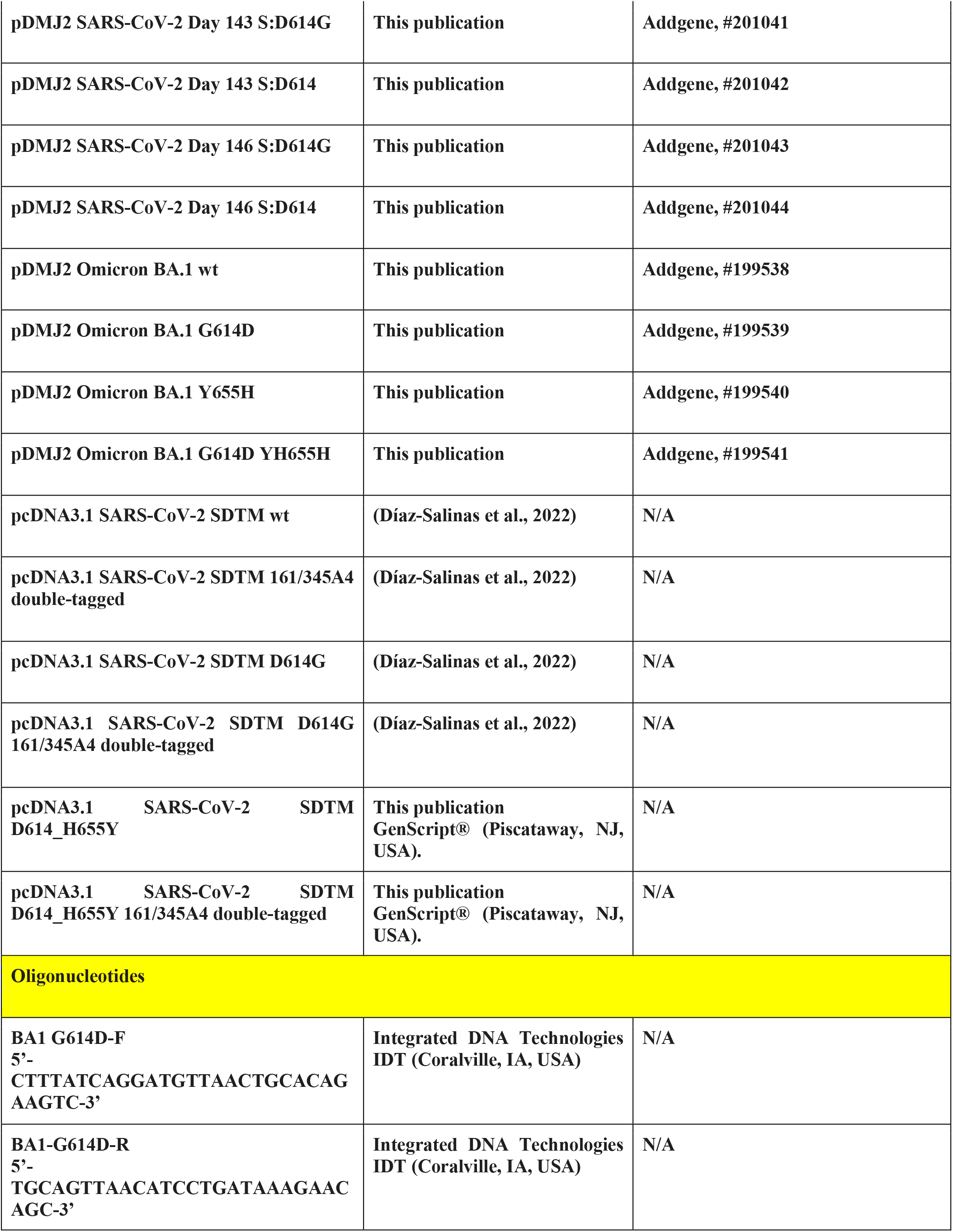
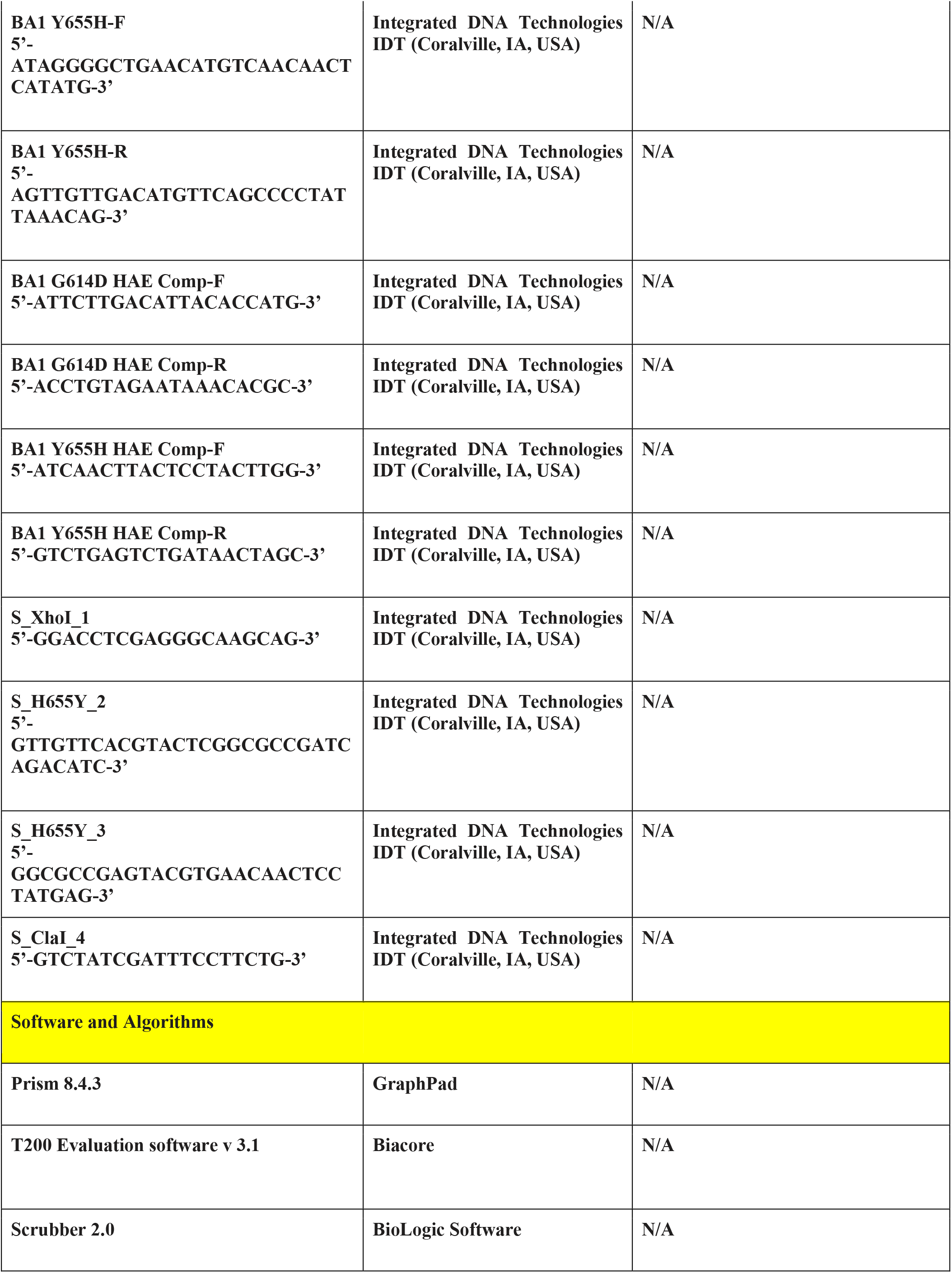
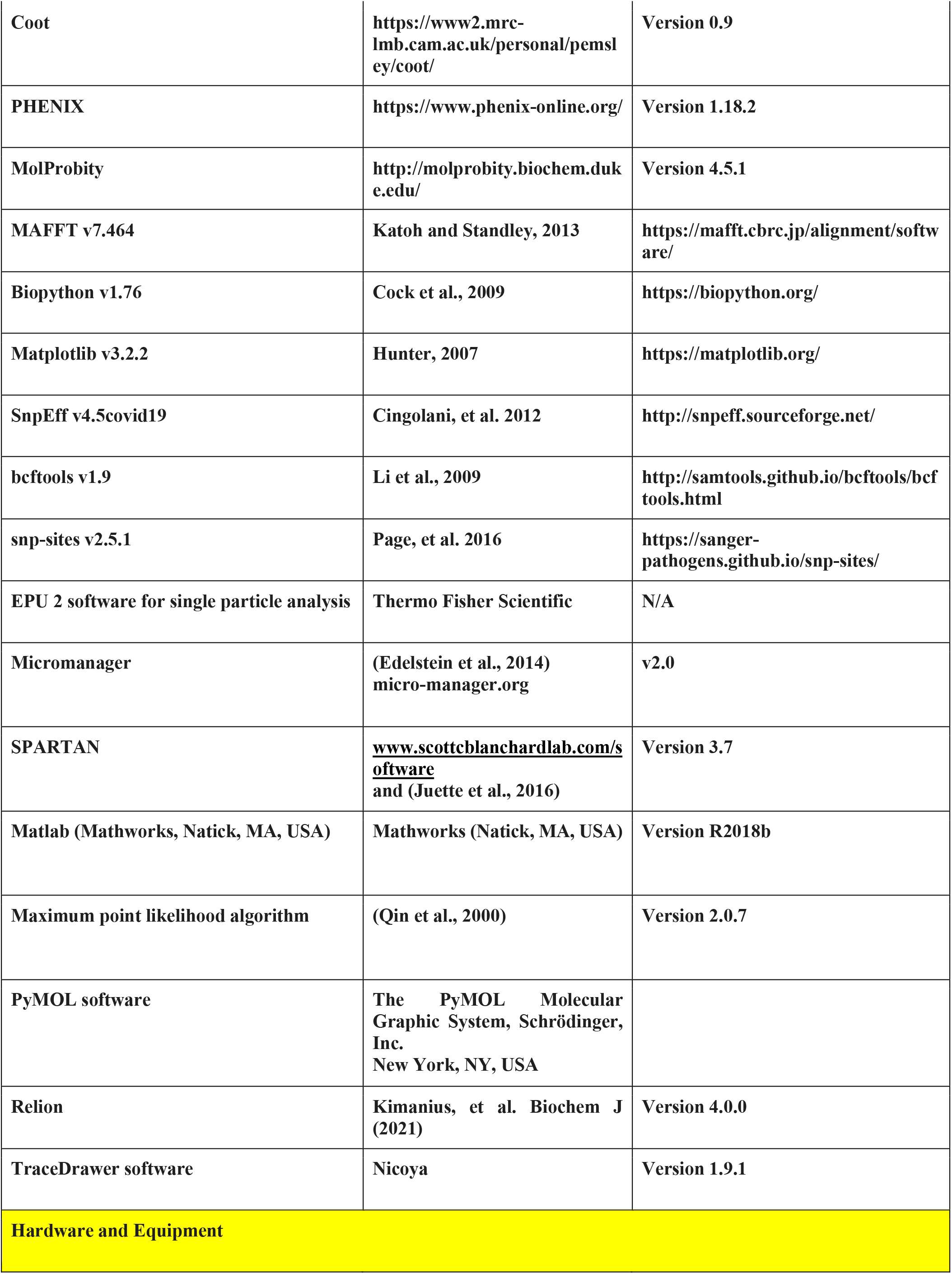
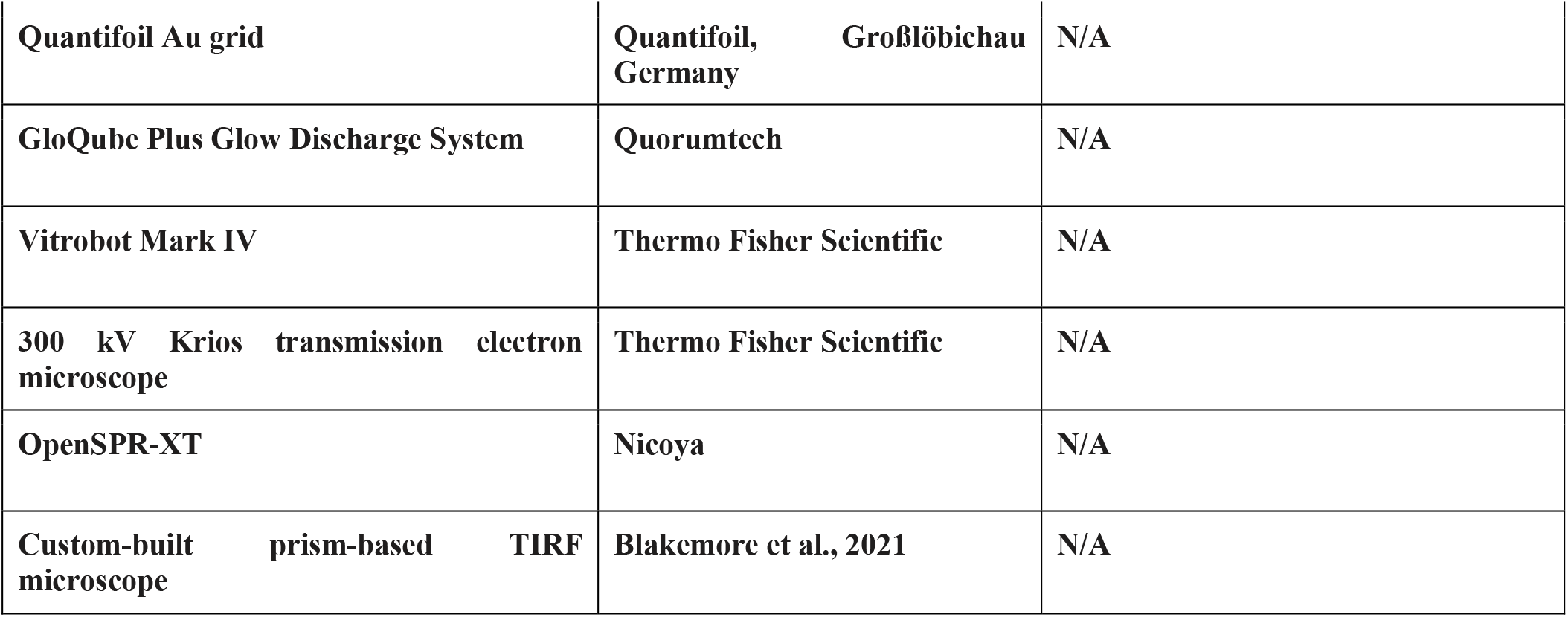
Reagents and Tools.

